# Comparative analysis of drug-like EP300/CREBBP acetyltransferase inhibitors

**DOI:** 10.1101/2023.05.15.540887

**Authors:** McKenna C. Crawford, Deepika R. Tripu, Samuel A. Barritt, Yihang Jing, Diamond Gallimore, Stephen C. Kales, Natarajan V. Bhanu, Ying Xiong, Yuhong Fang, Kamaria A. T. Butler, Christopher A. LeClair, Nathan P. Coussens, Anton Simeonov, Benjamin A. Garcia, Christian C. Dibble, Jordan L. Meier

## Abstract

The human acetyltransferase paralogs EP300 and CREBBP are master regulators of lysine acetylation whose activity has been implicated in various cancers. In the half-decade since the first drug-like inhibitors of these proteins were reported, three unique molecular scaffolds have taken precedent: an indane spiro-oxazolidinedione (A-485), a spiro-hydantoin (iP300w), and an aminopyridine (CPI-1612). Despite increasing use of these molecules to study lysine acetylation, the dearth of data regarding their relative biochemical and biological potencies makes their application as chemical probes a challenge. To address this gap, here we present a comparative study of drug-like EP300/CREBBP acetyltransferase inhibitors. First, we determine the biochemical and biological potencies of A-485, iP300w, and CPI-1612, highlighting the increased potency of the latter two compounds at physiological acetyl-CoA concentrations. Cellular evaluation shows that inhibition of histone acetylation and cell growth closely aligns with the biochemical potencies of these molecules, consistent with an on-target mechanism. Finally, we demonstrate the utility of comparative pharmacology by using it to investigate the hypothesis that increased CoA synthesis caused by knockout of PANK4 can competitively antagonize binding of EP300/CREBBP inhibitors and demonstrate proof-of-concept photorelease of a potent inhibitor molecule. Overall, our study demonstrates how knowledge of relative inhibitor potency can guide the study of EP300/CREBBP-dependent mechanisms and suggests new approaches to target delivery, thus broadening the therapeutic window of these preclinical epigenetic drug candidates.

## Introduction

Lysine acetylation is a prevalent posttranslational modification (PTM) whose therapeutic targeting represents an emerging paradigm in oncology.^1^ Foremost amongst the human enzymes that catalyze this modification are the multidomain protein paralogues EP300 and CREBBP, which due to their highly similar acetyltransferase active sites (>95% identity) are commonly referred to as EP300/CREBBP (Fig. 1).^2^ The first characterized substrates of EP300/CREBBP were histone proteins.^3, 4^ Canonically, EP300/CREBBP-catalyzed acetylation of histone H3 on lysines 18 and 27 (H3K18Ac and H3K27Ac) is associated with transcriptional activation. More recently, proteome-wide analyses have also implicated EP300/CREBBP as a major driver of highly penetrant lysine acetylation events in non-histone proteins.^5^ Furthermore, biochemical and cellular evidence supports the ability of EP300/CREBBP to catalyze non-canonical lysine acylations via the utilization of alternative metabolic cofactors such as propionyl-, butyrl-, and crotonyl-CoA, expanding their functional repertoire.^6^ Due to the critical role of acetylation in transformation and oncogenic transcription, EP300 and CREBBP have emerged as cancer drug targets.^7, 8^ However, these enzymes are also required for normal mammalian cell growth and development.^9^ Therefore, the therapeutic window for targeting EP300/CREBBP in advanced malignancies remains a topic of interest and active investigation.

**Figure 1.**
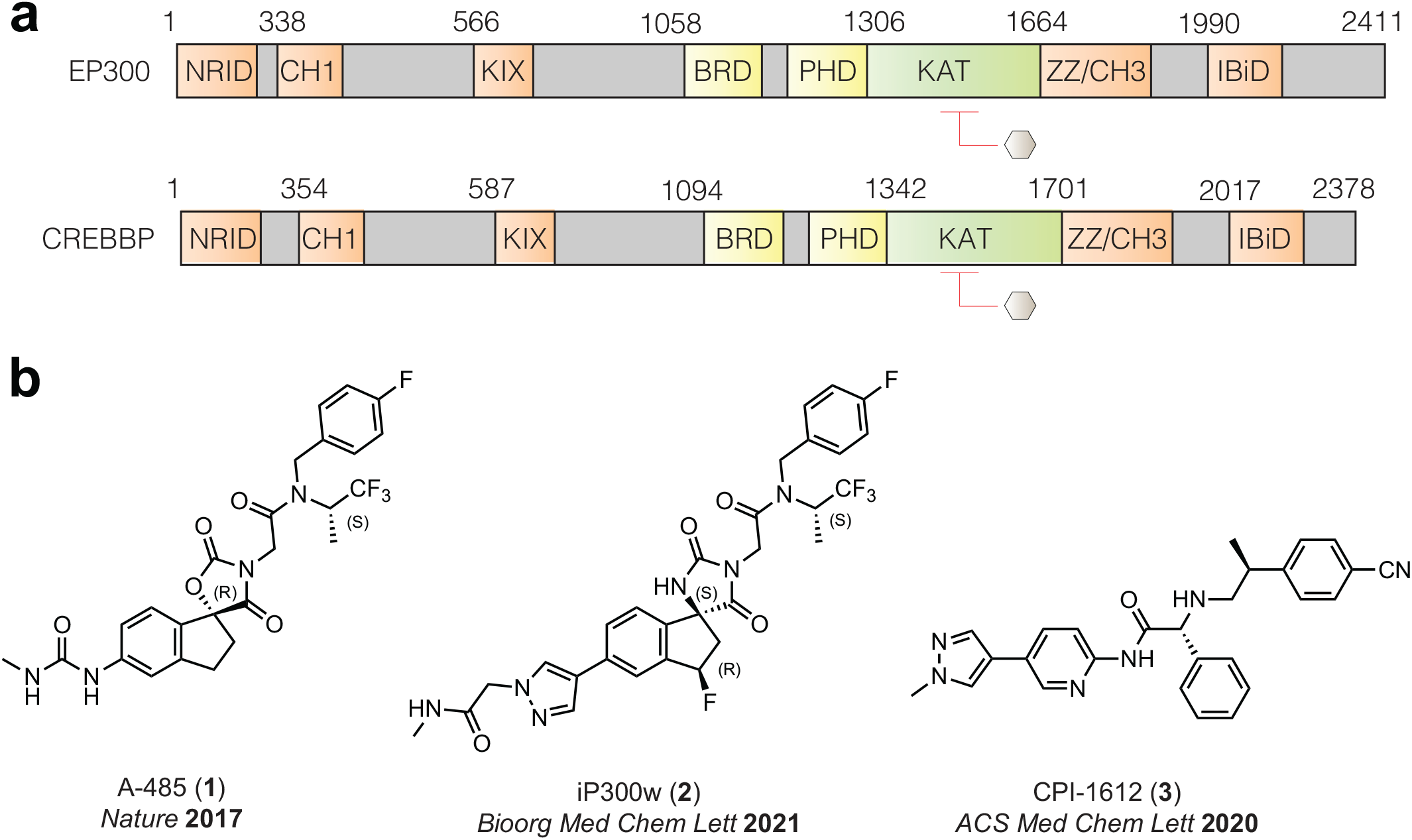
(a) Domain architecture of EP300 and CREBBP transcriptional coactivators. (b) Structure and primary reference for drug-like acetyltransferase inhibitors **1-3** comparatively assessed in this study.

To better understand the role of EP300/CREBBP in biology and disease, substantial work has been devoted to developing small molecule inhibitors of their acetyltransferase domains.^10^ The spiro-oxazolidinedione A-485 (**1**) was the first drug-like EP300/CREBBP acetyltransferase inhibitor to become commercially available and currently constitutes the most widely applied probe in the literature (**Fig. 1b**).^11^ Further medicinal chemistry optimization of this scaffold led to spiro-hydantoin iP300w (**2**), which differs from A-485 (**1**) by several heteroatom replacements, as well as an inverted stereocenter.^12^ Both the chiral and racemic forms of **2** have been applied in several studies.^13-16^ CPI-1612 (**3**) (**Fig. 1b**) is a structurally distinct aminopyridine EP300/CREBBP inhibitor developed in parallel to **1** and **2** optimizing for potency, and in vivo bioavailability but has been applied in a relatively small number of studies.^17^ Given that acetyltransferases were long considered an “undruggable” epigenetic enzyme family, it is remarkable that the field now has three drug-like compounds that can be used to probe EP300/CREBBP-catalyzed acetylation. However, this pharmacological windfall creates an urgent need to understand the relative properties of these molecules for appropriate and effective deployment. Specifically, the selectivity of these inhibitors for EP300/CREBBP are reported but their relative potencies are difficult to assess. While it has been experimentally established that these compounds compete with acetyl-CoA, their reported half-maximal inhibition constant (IC_50_) values are highly dependent on the acetyl-CoA concentration used in the biochemical reaction, which varies across studies.^11, 12, 17^ Furthermore, to our knowledge, these compounds have never been assessed in a common cellular model. To address this, here we report herein a comparative biochemical and cellular assessment of drug-like EP300/CREBBP inhibitors. First, we directly compared the relative inhibitory properties of **1-3**, discovering they possess distinct IC_50_ values and range broadly in terms of potency at physiological concentrations of acetyl-CoA. Next, we performed cellular studies, which revealed the ability of these compounds to inhibit histone acetylation and cell growth directly correlating with their biochemical potency. Finally, this knowledge of the comparative properties of **1-3** enabled two applications: 1) testing the hypothesis that increases in acetyl-CoA caused by knockout of PANK4 can alter the efficacy of EP300/CREBBP inhibitors in a human mammary epithelial cell line, and 2) demonstrating that use of potent EP300/CREBBP inhibitor scaffolds can enable stimuli-responsive inhibition as a proof-of-concept for targeted delivery efforts. Overall, our study reveals how knowledge of relative inhibitor potency can guide the study of EP300/CREBBP-dependent mechanisms while also suggesting targeted delivery of potent EP300/CREBBP inhibitors can broaden the therapeutic window of these pre-clinical epigenetic drug candidates.

## Results

### Biochemical assessment of EP300/CREBBP inhibition in the presence of physiological acetyl-CoA

A unifying aspect of EP300/CREBBP probes **1-3** is they are all acetyl-CoA competitive, which causes their inhibition to scale linearly with the amount of cofactor present in the enzyme reaction. Quantitative metabolomic and imaging studies have estimated the whole-cell concentration of acetyl-CoA to reside near the 5-15 micromolar range.^18-22^ However, IC_50_ values for EP300/CREBBP inhibitors have typically been determined at much low (nanomolar) concentrations of acetyl-CoA.^11^ To enable comparison of the biochemical properties of these three inhibitors, we developed a time-resolved fluorescence resonance energy transfer (TR-FRET) assay to assess the ability of **1-3** to disrupt activity of the EP300 catalytic domain (**Fig. 2a**).^11^ Compounds **1-3** were obtained from commercial suppliers and tested for their ability to inhibit acetylation of a biotinylated histone H3 peptide (human; amino acids 1-21) upon incubation with EP300 and acetyl-CoA in assay buffer. Reaction mixtures were incubated for 1 h prior to addition of Alexa Fluor 647-labeled anti-H3K9Ac antibody and Europium-labeled streptavidin. Determination of kinetic parameters in the presence of varying amounts of peptide substrate and acetyl-CoA cofactor yielded Michaelis constants (K_m_) of 18.2 nM and 73 nM, respectively (**Fig. S1**). These results prompted us to initially perform the assay at 50 nM, approximating the K_m_ for acetyl-CoA. Comparing IC_50_ values obtained under these conditions, CPI-1612 (**3**) emerged as the most potent EP300/CREBBP acetyltransferase inhibitor (IC_50_ = 10.7 nM), followed by iP300w (**2**) (IC_50_ = 15.8 nM) and A-485 (**1**) (IC_50_ = 44.8 nM). High concentrations of acetyl-CoA (5 μM, ∼100x > K_m_) reduced the inhibitory potency of each compound tested while maintaining the same trends (**Fig. 2b-c**). At this higher level of acetyl-CoA – which approximates the lower range of whole-cell metabolomic measurements -- inhibition of EP300/CREBBP by A-485 (**1**) occurs in the micromolar range (IC_50_ = 1.3 μM). This is similar to the concentration of A-485 required to observe reductions in histone acetylation in several cellular studies (**Fig. 2c**).^11, 23, 24^ Additionally, it was observed that A-485 (**1**) exhibited a greater shift in IC_50_ (∼29x) at higher acetyl-CoA concentrations compared to either iP300w (**2**) (∼7x) or CPI-1612 (**3**) (∼2x). This may be a reflection of assay conditions where inhibitors are pre-incubated with EP300 prior to addition of acetyl-CoA enabling compounds with low off-rates to maintain inhibition at high concentrations of cofactor.^25^ These studies establish a range of cofactor-dependent biochemical potencies for drug-like inhibitors of EP300/CREBBP.

**Figure 2.**
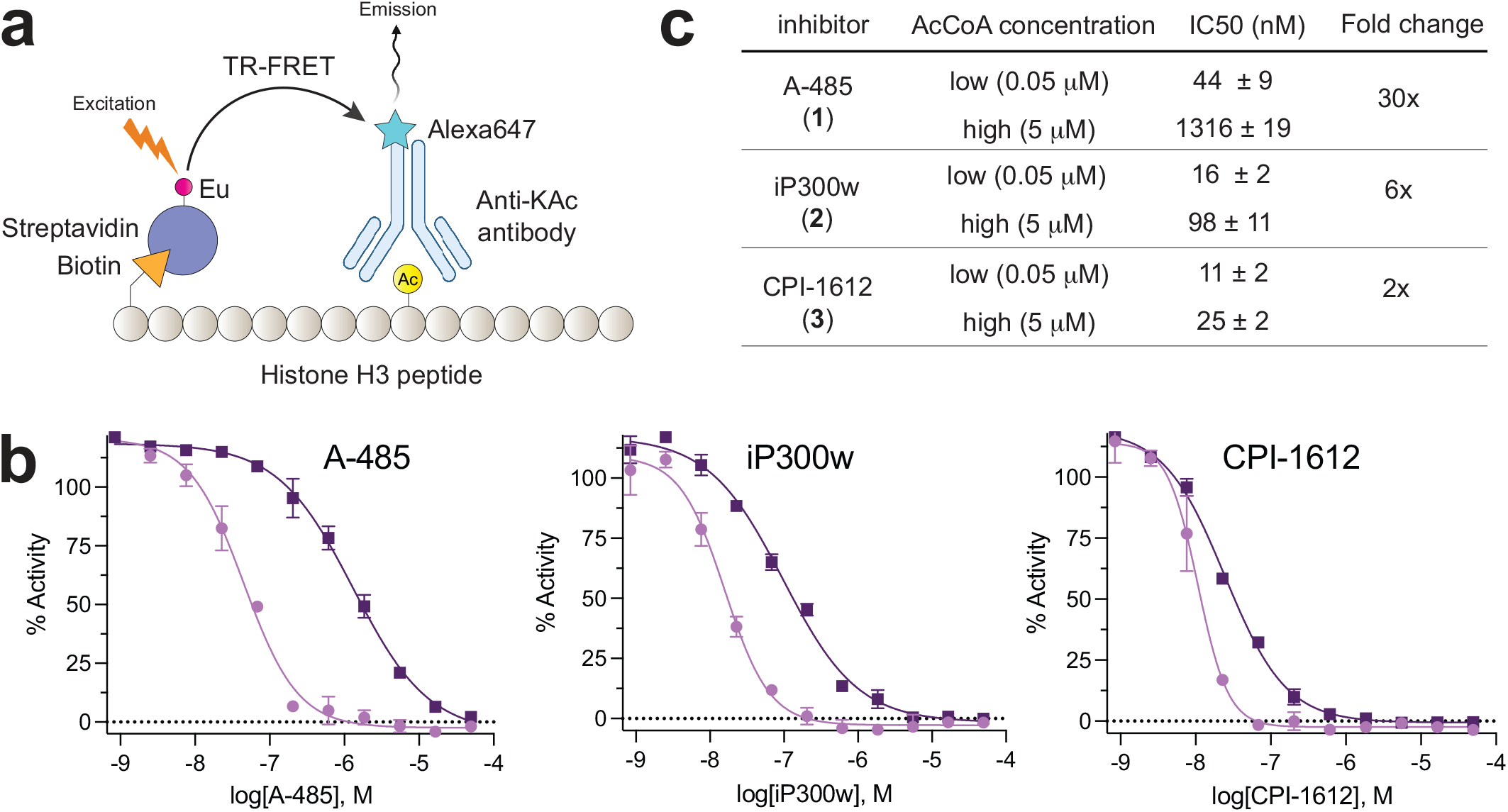
(a) TR-FRET assay for EP300-catalyzed acetylation of a histone H3 peptide. (b) Biochemical inhibition of EP300 by drug-like acetyltransferase inhibitors **1-3**. (c) Tabulated half-maximal inhibition concentrations for inhibitors. Fold-change represents the IC_50_ at high acetyl-CoA concentrations relative to the IC_50_ at high acetyl-CoA concentrations. Values represent the average of n=2 technical replicates.

### Comparative cellular assessment of EP300/CREBBP inhibitors

Next, we sought to quantitatively benchmark the properties of **1-3** in living cells. EP300 and CREBBP modify a wide variety of substrates in biochemical experiments and upon ectopic overexpression.^2^ However, two well-characterized cellular histone lysine substrates of these enzymes are H3K18 and H3K27.^26^ Even though H3K18Ac is the more abundant of the two markers, H3K27Ac is also of great interest due to its association with active enhancer regions. As a model to compare the effects of **1-3**, we chose MCF-7 breast cancer cells, which have previously been shown to respond to EP300/CREBBP inhibition.^27^ Cells were treated with escalating doses of **1-3** and harvested at 3 h to avoid overt cytotoxicity and allow the direct effects on acetylation to be observed (**Fig. 3a**). In a similar trend to our biochemical observations, CPI-1612 (**3**) showed the most potent activity when assayed across equivalent concentrations (8 nM – 5 μM), followed by iP300w (**2**) and A-485 (**1**) (**Fig. 3b**). Control experiments indicated levels of H3K23Ac, which is not regulated by EP300/CREBBP, were not affected.

**Figure 3.**
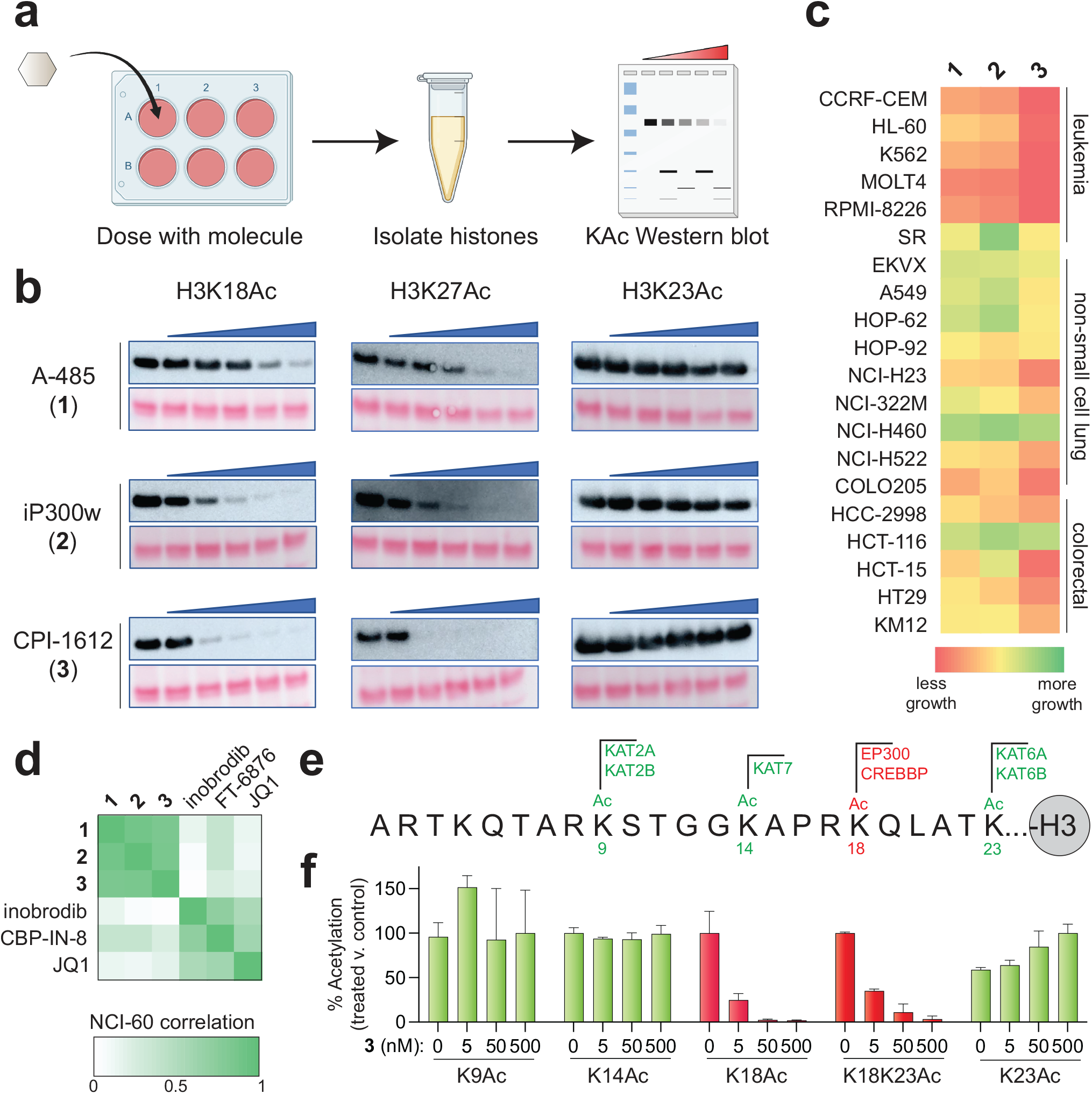
(a) Schematic for cell-based analysis of **1-3**. (b) Western blot analysis of histones extracted from MCF-7 cells after treatment with **1-3**. The left lane of each gel corresponds to vehicle-treated control, followed by cells incubated with five-fold increasing concentrations of **1, 2**, or **3** (8, 40, 200, 1000, 5000 nM). Exposure times are the same within each histone acetylation mark. (c) Heat-map depiction of growth caused by treatment of cells with **1-3** in the NCI-60 cell line screen. (d) Cross correlation analysis of NCI-60 inhibition by **1-3** and bromodomain inhibitors (e) Sequence of histone H3 N-terminus indicating sites of acetylation and the KAT responsible for modification. EP300/CREBBP-regulated K18Ac is colored in red. (f) Quantitative LC-MS analysis of histone H3 acetylation. Values correspond to the normalized intensities observed for histone peptides containing the specified modifications isolated from cells treated with **3**, with signal for untreated cells set equal to 100%. The intensity of singly modified peptides was used, in all cases with the exception of K14Ac, whose value was derived from the summed intensity of singly and doubly modified K14Ac peptides (2^nd^ modification: K9me1, K9me2, or K9me3). Values represent the average of n=3 biological replicates.

In principle the phenotype elicited by any single small molecule can be due to either an on- or off-target effect; however, if multiple chemically distinct small molecules designed to target a single gene all elicit the same phenotype, one’s confidence in their on-target mechanism becomes greater. As such, we sought to apply our knowledge of the comparative potencies of **1-3** to test whether the simplest phenotype caused by EP300/CREBBP inhibitors – cell death – was consistent with their effects on histone acetylation and a putative on-target mechanism. Compounds **1-3** were submitted for analysis in the NCI-60 screen, which has been used since 1990 to analyze the growth inhibitor effects of small molecules across a range of transformed cell lines derived from diverse tissues of origin including colorectal, renal, ovarian, prostate, lung, breast, and central nervous system tumors.^28^ Consistent with our biochemical and cellular acetylation assays, CPI-1612 (**3**) again emerged as the most potent inhibitor of cell growth followed by iP300w (**2**) and A-485 (**1**) (**Figs. 3c and Table S1**). These trends were further confirmed using an orthogonal ATP-Glo assay in MCF-7 cells (**Fig. S2**). Subsequently, the COMPARE algorithm was used to compare the patterns of growth inhibition elicited by **1-3** in the NCI-60 assay against the >150,000 compounds analyzed in the screen to date.^29^ This analysis revealed that **1-3** most closely cluster with each other consistent with engagement of a common target, presumably EP300/CREBBP (**Fig. 3d**). Interestingly, known inhibitors of both EP300/CREBBP^30, 31^ and BET family^32^ bromodomains were subjected to COMPARE analysis and were found to be less correlated with **1-3**, suggesting these two classes of compounds could serve as distinct modulators of acetylation-dependent signaling cascades (**Figs. 3d, S3, and Table S1**). Finally, the specificity of CPI-1612 (**3**) was directly assessed using quantitative LC-MS analysis of histone acetylation in treated cells,^33^ focusing on four abundant histone acetylation markers whose writers are well-characterized: H3K9Ac (written by KAT2A/B), H3K14Ac (KAT7), H3K18Ac (EP300/CREBBP), and H3K23Ac (KAT6A/B; **Fig. 3e**).^26^ This revealed CPI-1612 (**3**) affected the abundance of only the EP300/CREBBP-dependent peptides containing H3K18Ac (**Fig. 3f and Table S2**). An increase in the levels of singly modified H3K23Ac peptide coincided with specific loss of H3K18Ac from K18/K23 diacetylated peptide, indicative of orthogonal regulation. Overall, the parallel trends in the biochemical and cellular potencies of EP300/CREBBP inhibitors are consistent with an on-target mechanism for **1-3** and further highlight the relatively high potency of CPI-1612 (**3**).

### Comparative pharmacology enables probing of metabolically regulated EP300/CREBBP inhibition

Recently, we reported the discovery of a phosphorylation-dependent signaling cascade that can upregulate CoA biosynthesis in human cells by inhibiting the activity of PANK4, a negative regulator of this pathway (**Fig. 4a**).^34^ This occurs via PANK4’s ability to reverse the rate-limiting step in CoA production, which is phosphorylation of pantothenate (vitamin B5) by PANK1-3.^35^ Removing this hurdle causes PANK4-deficient cells to harbor constitutively elevated concentrations of acetyl-CoA. In independent studies, Bishop et al. reported that long-term exposure of multiple myeloma cells to A-485 (**1**) can select for clones with activating mutations in PANK3 or inactivating mutations in PANK4 that upregulate CoA biosynthesis.^36^ The most parsimonious explanation for this finding is that overproduction of acetyl-CoA decreases the efficacy of EP300/CREBBP inhibitors by directly competing for active site occupancy, as observed in our biochemical experiments (**Fig. 2b-c**). However, acetyl-CoA plays a central role in a wide-range of biological processes including lipid signaling, cholesterol biosynthesis, and protein homeostasis.^37^

**Figure 4.**
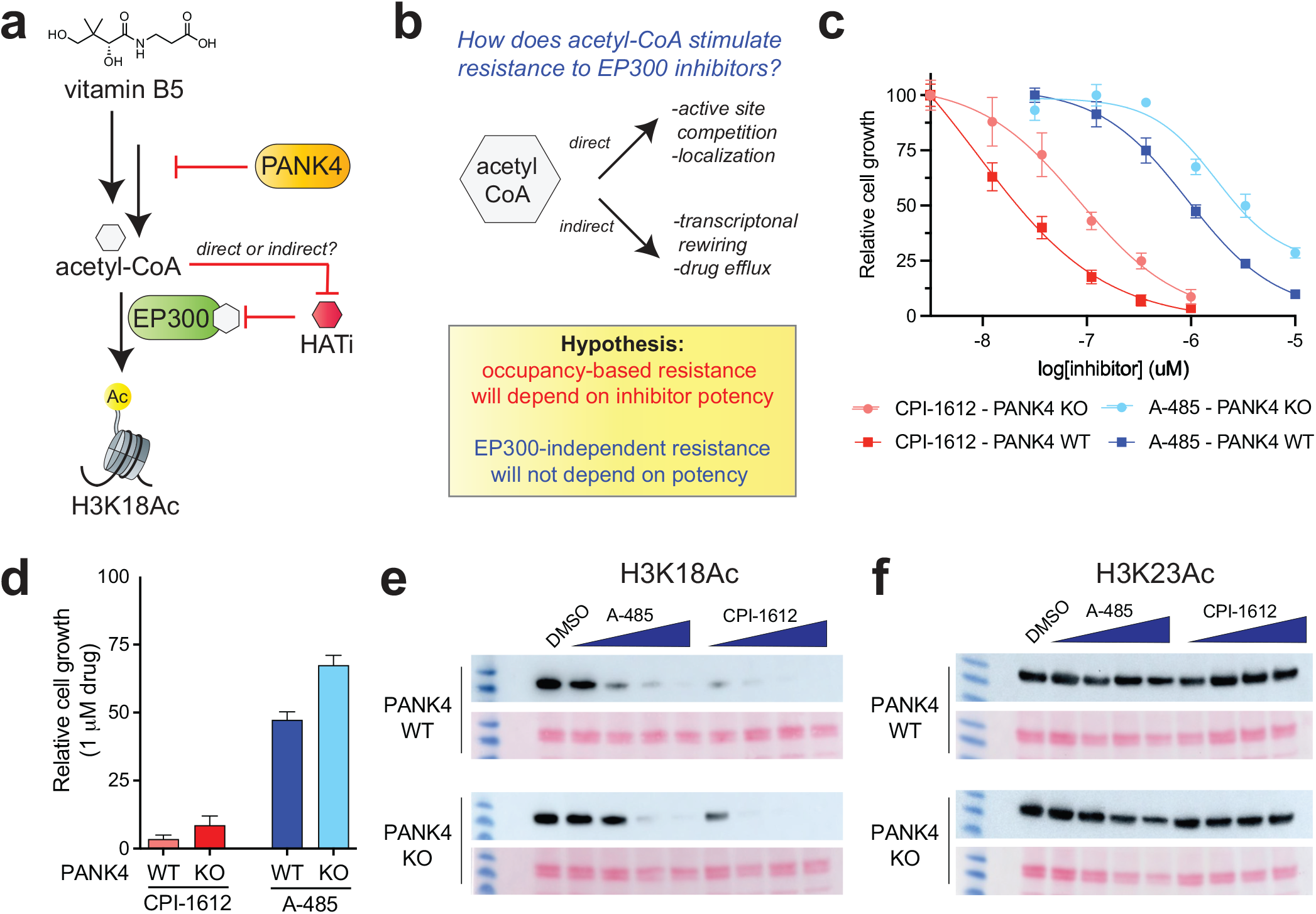
(a) PANK4 inhibits the production of acetyl-CoA from vitamin B5 by consuming a metabolic intermediate (4’-phosphopantetheine, not shown). Knockout of PANK4 can increase acetyl-CoA levels and cause resistance to inhibitors of EP300. (b) Rationale for how inhibitors with distinct potencies can be used to differentiate direct and indirect acetyl-CoA-dependent resistance mechanisms. (c) Dose-response curves of MCF-10A cells (PANK4 KO+WT and PANK4 KO+EV) grown in the presence of **1** or **3**. (d) Relative proliferation of MCF-10A cells (PANK4 KO+WT and KO+EV) grown in the presence of 1 μM **1** or **3**. (e) Effects of 3 h treatment of **1** and **3** ((8, 40, 200, 1000, 5000 nM) on H3K18Ac and (f) H3K23Ac in MCF-10A cells (PANK4 KO+WT and PANK4 KO+EV). Exposure times are the same within each histone acetylation mark.

Conceivably, rewiring of cellular pathways by high acetyl-CoA could decrease the efficacy of EP300/CREBBP inhibitors indirectly, by promoting drug efflux or EP300-independent proliferation. Considering how to resolve these mechanisms, we wondered if this may be an area where knowledge of the comparative pharmacology of EP300/CREBBP inhibitors could be usefully leveraged. Specifically, we hypothesized that if PANK4 inactivation (knockout; KO) led to a resistance mechanism dependent on EP300/CREBBP active site occupancy (e.g. competitive binding), then the differential potency of **1** and **3** would be preserved – albeit over a shifted concentration range – in the resistant PANK4 KO cell line (**Fig. 4b**). In contrast, if PANK4 inactivation led to a resistance mechanism independent of EP300/CREBBP active site occupancy (e.g., drug efflux), then the differential potency of **1** and **3** would be lost. To test this, we examined the effects of EP300/CREBBP inhibitors in non-transformed MCF10A mammary epithelial cells in which PANK4 had been knocked out and either a wild-type PANK4 gene (WT) or an empty vector (EV) control had been stably re-expressed.^34^ Consistent with our previous observations of PANK4-dependent acetyl-CoA and the recent results of Bishop et al.,^36^ PANK4 KO cells expressing EV displayed approximately 2-8-fold decreased sensitivity to EP300/CREBBP inhibition compared to cells in which the WT PANK4 had been re-introduced (**Fig 4c**). However, the relative trends in growth inhibition between inhibitors was maintained. For example, at a concentration of 1 μM A-485 (**1**) is nearly inactive in PANK4 KO+EV cells (**Fig. 4c-d**). In contrast, CPI-1612 (**3**) remains active against both PANK4 KO+WT and KO+EV cells at this concentration but shows a similar reduction in efficacy in KO+EV cells as its dosage is decreased to 100 nM, consistent with an occupancy-based effect (**Fig. 4d**). KO cells transformed with empty vector (EV) and catalytically inactive PANK4 mutant (KO+mutant) behaved similarly (**Fig. S4**). To correlate these observations with the effect of the compounds on cellular acetyltransferase activity, we monitored histone H3 acetylation at an EP300/CREBBP-dependent site (H3K18Ac) and EP300/CREBBP-independent site (H3K23Ac). This revealed nearly identical trends, with PANK4 KO lessening the effects of both **1** and **3** at micromolar and nanomolar concentrations, respectively (**Fig. 4e**). H3K23Ac did not significantly respond to inhibitor treatment in either cell line (**Fig. 4f**). These results indicate that the PANK4-dependent activity of EP300/CREBBP inhibitors can extend beyond multiple myeloma models and likely acts via an occupancy-based mechanism, potentially by causing a significant change in the concentration of acetyl-CoA accessible to EP300/CREBBP. At equimolar concentrations, CPI-1612 (**3**) is less susceptible to this mechanism compared to A-485 (**1**) (**Fig. 4d**). The ability of this mechanism to be selected for in vivo during cancer treatment, or in the presence of more potent EP300/CREBBP inhibitors such as **2-3**, remains unknown.

### Comparative pharmacology enables stimuli-sensitive EP300/CREBBP inhibitor release

One potential reason EP300/CREBBP acetyltransferase inhibitors are not actively being evaluated in clinical trials is on-target toxicity.^10^ Thus, these molecules may benefit from strategies to direct them more specifically to malignant – as opposed to healthy – tissues. Light-activated drug release provides one potential approach.^38^ Due to the non-quantitative nature of most photouncaging reactions, these methods are most effective when unleashing potent payloads that can occupy the intended target at low concentrations of drug.^39^ Considering the studies above we wondered: could the high potency of the CPI-1612 scaffold enable conditional release of an active EP300/CREBBP inhibitor? To explore this, we prepared a photocaged CPI-1612 analogue (**4**) (**Fig. 5a**). The design of this molecule was based on structural data, which suggests derivatization of CPI-1612’s secondary amine with a bulky group should significantly impede EP300/CREBBP binding (**Fig. 5b**),^17^ as well as patent literature, which reports the product of photocleavage (**5**) has potent in vivo antitumor activity.^40^ Parent inhibitor **5** was synthesized based on the published route^40^ and derivatized with nitrobenzyl chloroformate to produce the photocaged analogue **4** (**Scheme S1**). Consistent with our expectation, biochemical characterization via TR-FRET assay indicated that nitrobenzyl modification of the secondary amine decreases the EP300/CREBBP inhibitor potency of **4** ∼50-fold relative to uncaged analogue **5** (**Fig. 5c**). HPLC reaction monitoring confirmed the facile conversion of **4** to **5** upon exposure to 302 nm or 365 nm light in aqueous solution or buffer (**Figs. 5d and S5**). To test whether active EP300/CREBBP inhibitor release could be controlled by light, we incubated cells with **5** (10 or 25 μM), washed to remove inhibitor in media, and irradiated them for 1 min at 302 nm (**Fig. 5e**). Cells were then incubated for 3 h, harvested, and assessed for changes in EP300/CREBBP-dependent H3K18Ac. This analysis revealed that cells that were both treated with **4** and exposed to light (UV) exhibited clear decreases in H3K18Ac (**Fig. 5c**). However, cells only treated with **4** or only irradiated showed no such effect. Importantly, the potency of the CPI-1612 scaffold is critical to this result as it enables a short burst of UV irradiation to uncage an active concentration of EP300/CREBBP inhibitor. Intuitively, this task would be expected to be more challenging with a less active compound such as A-485 (**1**). Although beyond the scope of the current study, we envision targeted release of potent EP300/CREBBP inhibitors could find utility in malignancies already treated using phototherapy, such as head and neck cancers,^41^ as well as valuable applications in basic science.^42^ Overall, our studies demonstrate how a knowledge of the pharmacology of EP300/CREBBP inhibitors can guide the design of new chemical biology tools and effective stimuli-dependent release strategies.

**Figure 5.**
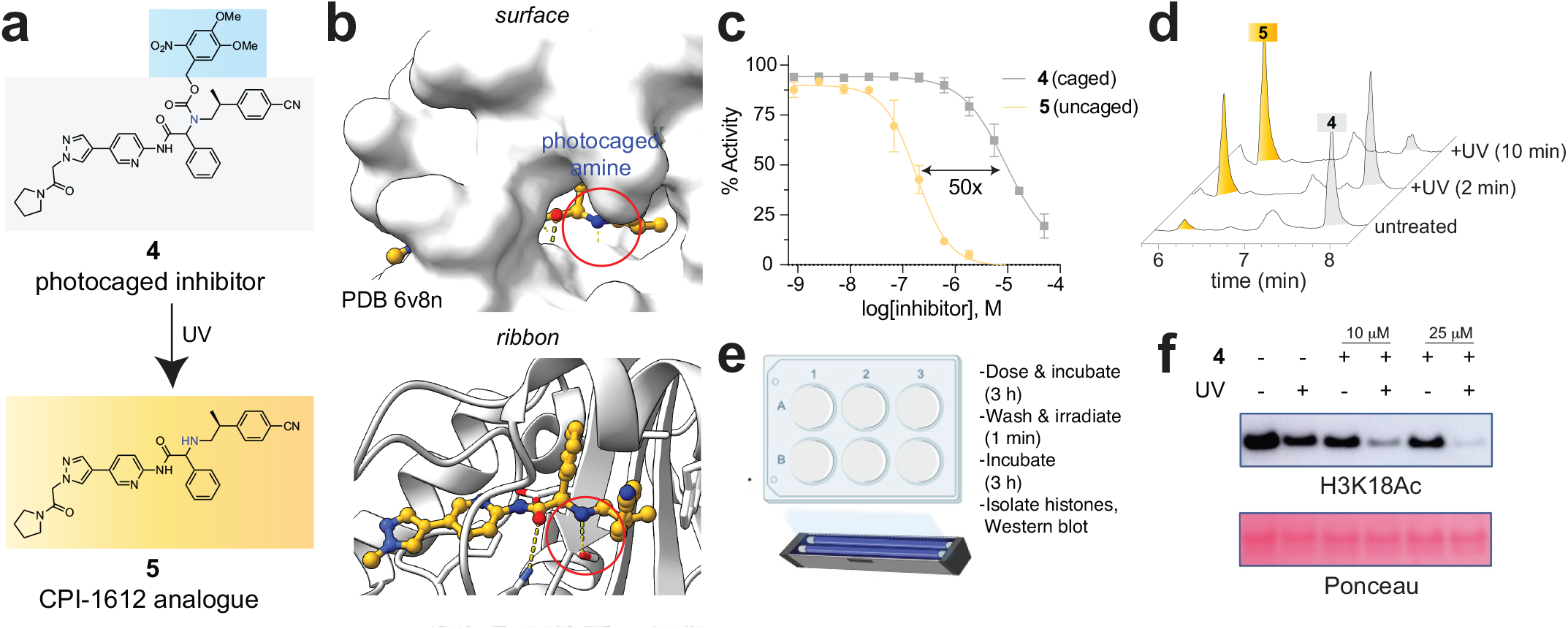
(a) Photocaged CPI-1612 analogue **4** and active inhibitor **5**. (b) Rationale for caging CPI-1612 analogues via derivatization of the secondary amine. (c) Relative EP300 biochemical inhibitor potency of **4** and **5** as assessed by TR-FRET histone acetylation assay (n=2 technical replicates). (d) Irradiation of **4** with UV light (302 nm) produces parent compound **5** in vitro. (e) Schematic for cellular photouncaging studies. (f) Light-dependent inhibition of EP300/CREBBP-dependent histone acetylation in living MCF-7 cells.

## Discussion

Here we report the comparative analysis of drug-like inhibitors of histone acetyltransferase paralogues EP300 and CREBBP. Our studies coalesce on four key findings:

1. The biochemical inhibitor potency of EP300/CREBBP inhibitors follows the order A-485 (**1**) < iP300w (**2**) < CPI-1612 (**3**). These trends are maintained in the presence of 5 μM acetyl-CoA, which lies near experimentally-determined measurements of whole-cell acetyl-CoA (∼5-15 μM).
2. The effects of **1-3** on cell growth and histone acetylation parallel their biochemical potency. These structurally distinct molecules also exhibit similar profiles of growth inhibition in the NCI-60 assay. CPI-1612 (**3**) also selectively modulates EP300/CREBBP-dependent acetylation in cellular histone H3 profiling assays. Together, our summed data strongly support a common mechanism for **1-3**, which we infer to be on-target inhibition of EP300/CREBBP.
3. The absolute potency of EP300/CREBBP inhibitors **1** and **3** are decreased by PANK4 KO, a genetic perturbation previously shown to increase the whole-cell concentration of acetyl-CoA. The relative potencies of **1** and **3** are maintained upon PANK4 KO, consistent with an occupancy-based resistance mechanism.
4. EP300/CREBBP inhibitors based on the CPI-1612 scaffold can be caged at the secondary amine to reduce their biochemical potency by approximately 50-fold. Photouncaging of aminopyridine **4** can release **5** at concentrations sufficient to modulate histone acetylation in living cells.

We anticipate these findings will be highly useful to researchers seeking to modulate EP300/CREBBP in living cells, in particular clarifying the context-dependent concentration regimens over which biochemical and cellular activity are observed. All available data indicate **1-3** are selective for the EP300/CREBBP acetyltransferase domain. We therefore suggest as a best practice for the field that **1-3** be used as lead probes to study EP300/CREBBP acetylation. A corollary is that employment of historic inhibitors – for whom selectivity data is ambiguous or lacking,^43^ and which continue to find prevalent use in the literature^44-48^ – should be avoided. The high potency exhibited by CPI-1612 (**3**) nominates it as an in vitro probe of choice and opens the door for applications where its potent activity may prove especially advantageous, including light and enzyme-mediated uncaging,^49^ antibody-drug conjugates,^50^ covalent probe design,^51^ and targeted protein degradation.^52, 53^ We envision the availability of well-characterized selective inhibitors of EP300/CREBBP will greatly benefit researchers wishing to study the role of protein acetylation in biology and disease.

## Acknowledgements

The authors thank Dr. Martin Schnermann (NCI) for helpful discussions and Jordan Nafie and Dr. Rina Dukor (BioTools, Inc.) for their assistance with the VCD analysis. This work was supported by the Intramural Research Programs of the National Cancer Institute, Center for Cancer Research ZIA BC011488 (J.L.M.) and the National Center for Advancing Translational Sciences, National Institutes of Health ZIA TR000327 (A.S.). This research was also supported by the V Foundation, V Scholar Grant V2019-009 (C.C.D), NIH R00-CA194314 (C.C.D), NIH F31-CA254169-01 (S.A.B.), and NIH CA196539 and HD106051 (B.A.G.). In addition, this project has been funded in part with federal funds from the National Cancer Institute, National Institutes of Health, under contract number HHSN261200800001E and HHSN261201500003I.

