## Supplementary Information for "Comparative analysis of drug-like EP300/CREBBP acetyltransferase inhibitors"

^1^Chemical Biology Laboratory, National Cancer Institute, Frederick, MD, USA. ^2^Department of Pathology, Cancer Research Institute, Beth Israel Deaconess Medical Center, Harvard Medical School, Boston, MA, USA. ^3^National Center for Advancing Translational Sciences, National Institutes of Health, Rockville, MD, USA. ^4^Department of Biochemistry and Molecular Biophysics, Washington University School of Medicine, St. Louis, MO, USA. ^5^Molecular Pharmacology Laboratories, Applied and Developmental Research Directorate., Frederick National Laboratory for Cancer Research, Frederick, MD, USA.

**Table of Contents for Supporting Information**

**Page**

Supplementary figures	S2

General materials and methods	S6

Biochemical assays	S7

Cytotoxicity analyses	S7

Histone modification analyses	S8

In vitro and cellular analysis of photocaged inhibitors	S11

Supplementary schemes	S12

Synthesis of photocaged inhibitor **4**	S14

Chiral separation and absolute configuration of **4** and **5**	S20

Full Western blot images	S24

References	S25

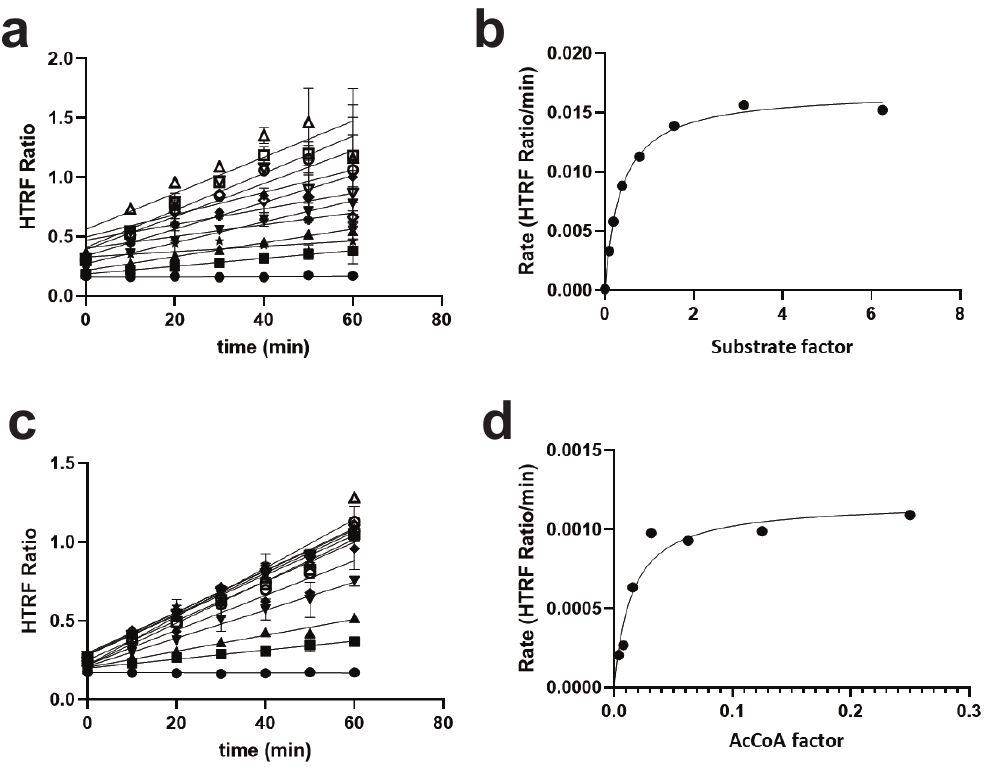

**Figure S1**. (a) Reaction progress curves for EP300 in the presence of varying histone H3 peptide and constant acetyl-CoA (50 nM). (b) Determination of Michaelis-Menten parameters for H3 1-21 peptide (K_m_ = 18.3 nM). (c) Reaction progress curves for EP300 in the presence of varying acetyl-CoA and constant histone H3 peptide (20 nM). (d) Determination of Michaelis-Menten parameters for acetyl-CoA (K_m_ = 73 nM).

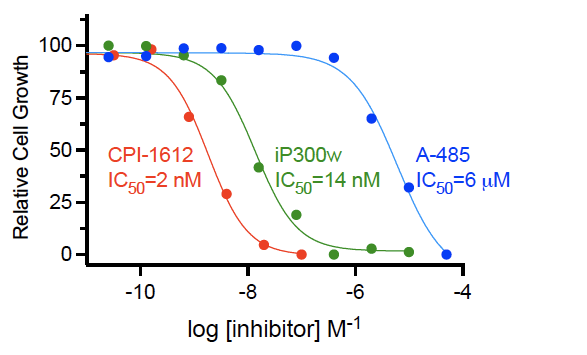

**Figure S2**. CellTiter-Glo-based determination of growth inhibition by **1**-**3** in MCF-7 cells.

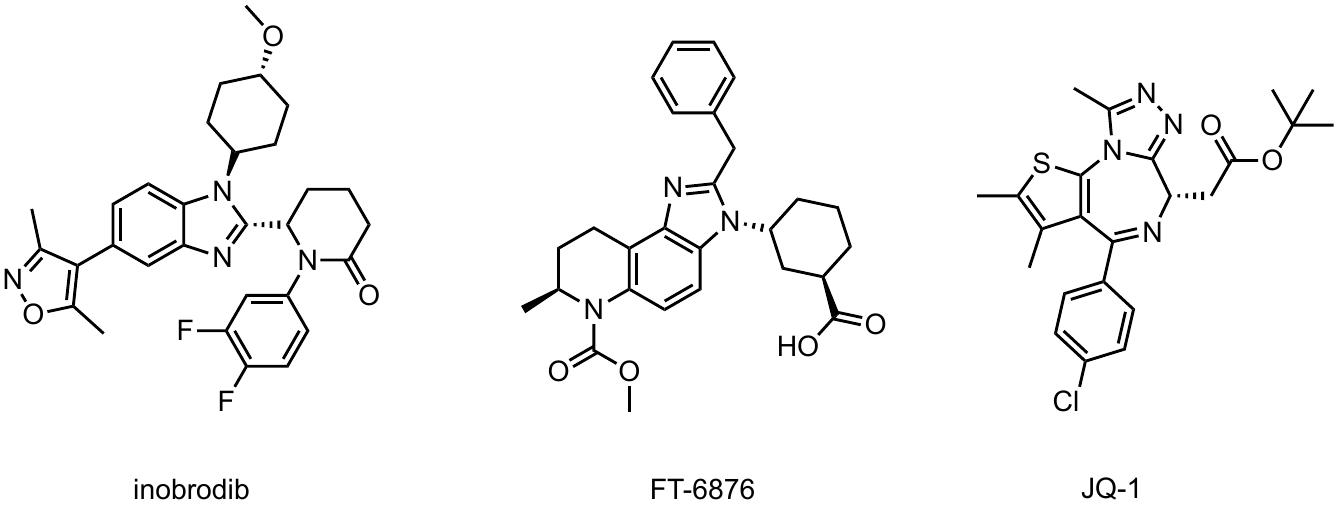

**Figure S3**. Structures of bromodomain inhibitors analyzed in Figure 3d.

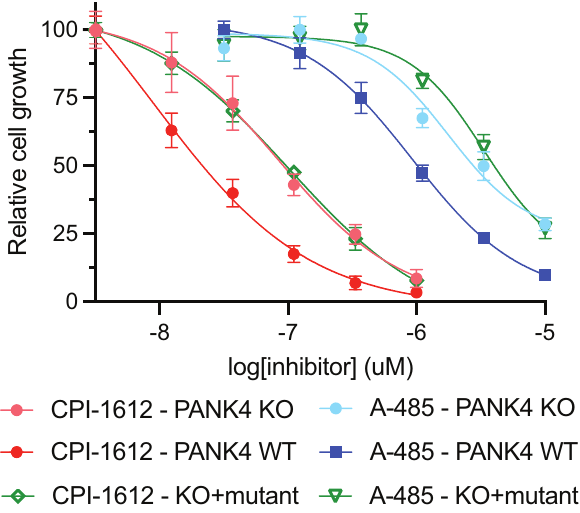

**Figure S4**. Dose-response curves of MCF-10A cells (KO+EV, KO+PANK4 WT and KO+PANK4 mutant) grown in the presence of **1** or **3**. IC_50_ values for A-485 **1**: 1760 nM (KO+EV), 929 nM (KO+PANK4 WT), 3370 nM (KO+PANK4 mutant). IC_50_ values for CPI-1612 **3**: 84 nM (KO+EV), 8.9 nM (KO+PANK4 WT), 102 nM (KO+PANK4 mutant).

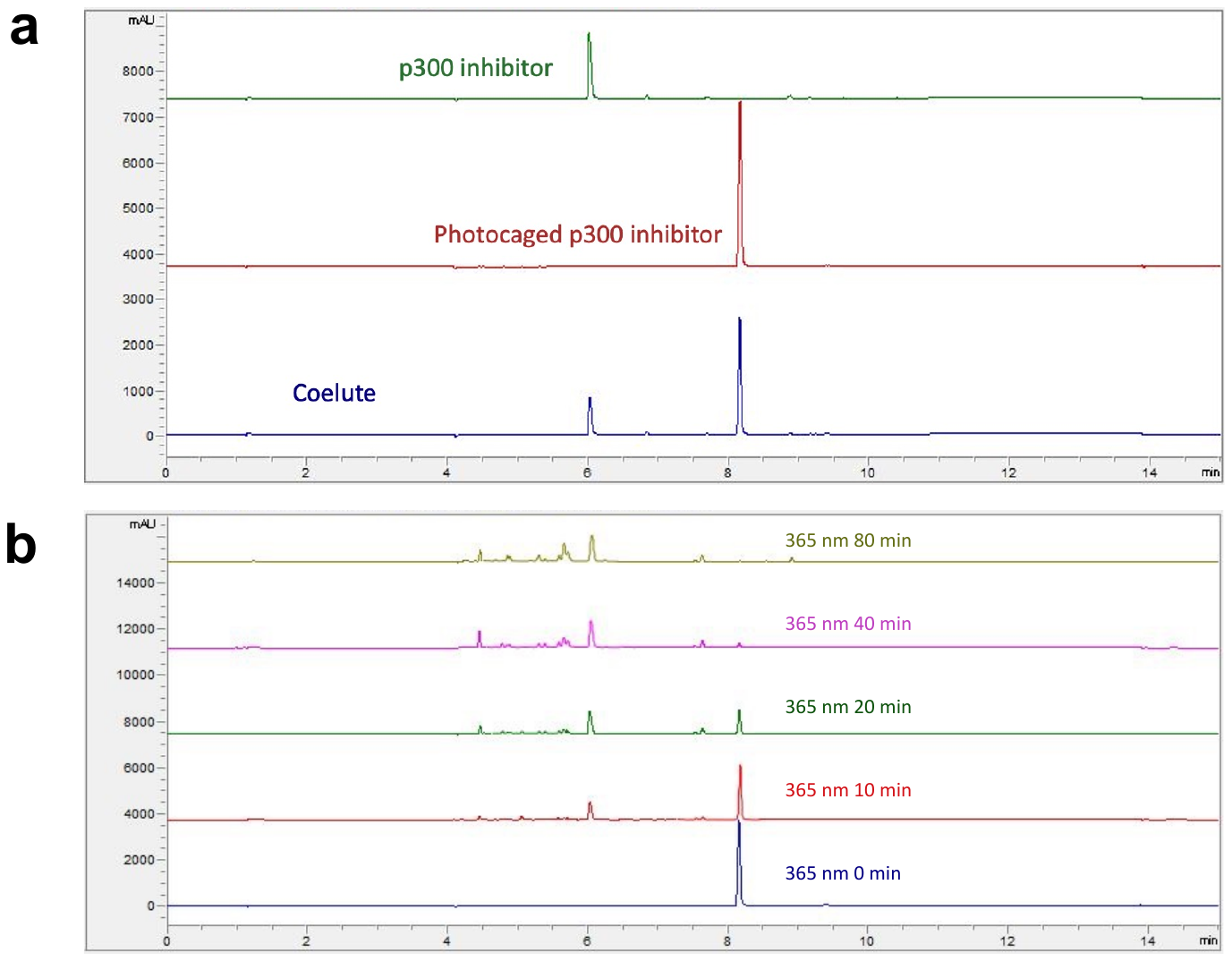

**Figure S5**. Photodeprotection of **4** upon irradiation with 365 nm light.

**General materials and methods**

Unless otherwise specified, all chemicals and solvents were purchased from Sigma, VWR, or Fisher and used without further purification. Analytical LC-MS of commercial EP300 inhibitors were carried out using an Agilent 1200 Quaternary LC-MS with an Agilent EC-C18 column (2.7 μM, 2.1 x 50 mm) employing a gradient of 0 → 30% acetonitrile/0.1% formic acid over 6 min at a flow rate of 0.6 mL/min. MCF-7 cells were cultured at 37 °C under 5% CO_2_ in EMEM (Quality Biological 112-016-101) with 10% FBS (Avantor Seradigm 97068-085), 2 mM L-glutamine (Quality Biological 118-084-721), and 0.01 mg/mL human recombinant insulin (MilliporeSigma 91077C). T-47D cells were cultured at 37 °C under 5% CO_2_ in RPMI-1640 (Quality Biological 112-024-101) with 10% FBS, 2 mM L-glutamine, and 0.2 u/mL human recombinant insulin. *PANK4* KO MCF10A cells (ATCC, CRL-10317) expressing empty vector, wild type PANK4, or a phosphatase-dead PANK4 point mutant (D623A)^1^ were maintained in standard MCF10A growth medium without antibiotics (DMEM/F12 medium (Wisent Bioproducts 319-075-CL), 5% horse serum (Gemini Bio 100508), 10 µg/mL insulin (Thermo Fisher Scientific/Gibco A11382II), 0.5 mg/mL hydrocortisone (Sigma-Aldrich H4001), 20 ng/mL EGF (R&D Systems, 236-EG-01M) and 100 ng/mL cholera toxin (List Biological Laboratories 100B)).

Precision Red Protein Assay (#ADV02) was purchased from Cytoskeleton. SDS-PAGE was performed using 4-12% Bis-Tris NuPAGE gels (Invitrogen #NP0322 and #NP0323), using XCell SureLock Mini-Cells (Invitrogen EI0002) and MES running buffer (Invitrogen #NP0002) following manufacturer’s protocols. BenchMark Pre-stained Protein Ladder was used for all gels (Invitrogen 10748010). For Western blotting, SDS-PAGE gels were transferred to 0.2 µM pore nitrocellulose membranes (Novex, Life Technologies # LC2000) by either wet transfer electroblotting at 30 volts for 1 h using a XCell II Blot Module (Invitrogen EI9051) and NuPAGE transfer buffer (Invitrogen NP00061) following the manufacturer’s protocols, or via iBlot dry transfer using the iBlot dry blotting system (Invitrogen IB1001) and nitrocellulose transfer stacks (Invitrogen IB301001) at 20 volts for 8 min following the manufacturer’s protocols. The specific transfer method used for each figure is specified in the full Western blot captions below. Total protein content on western blots was visualized by Ponceau staining followed by washing of membranes at least three times with 5% acetic acid in ddH_2_O. To visualize histone acetylation, membranes were blocked using StartingBlock (PBS) Blocking Buffer (Thermo Scientific 37538) for 30 min at room temperature and then incubated overnight at 4 °C in a solution containing the primary antibody at the indicated dilution. H3K18Ac (07-354) and H3K23Ac (07-355) were purchased from MilliporeSigma and used at 1:10,000 dilutions. H3K27Ac (8173S) antibody was purchased from Cell Signaling Technology and used at a 1:1,000 dilution. Membranes were washed the next day with 1x TBST at least 3 times and incubated for 1 h with anti-rabbit IgG HRP-linked antibody (Cell Signaling Technology 7074S) diluted to 1:1,000 in 1x TBST + 5% non-fat dry milk. Membranes were washed at least 3 times with 1x TBST, prior to imaging with Lumiglo (Cell Signaling Technology #7003) or SuperSignal ELISA Femto Substrate (Thermo Scientific 37074) according to manufacturer’s protocols. Colorimetric and chemiluminescent signals were detected using an Amersham ImageQuant 800 (Cytiva 29399482). Optical measurements were recorded on a Cytation 5 Multimode Plate Reader (Biotek) and PHERAstar FSX microplate reader with HTRF optical module (BMG LabTech).

**Biochemical assays**

A time-resolved fluorescence resonance energy transfer (TR-FRET) assay was used to assess the ability of **1**-**3** to disrupt the activity of EP300 catalytic domain. Briefly, an enzyme mix was made consisting of EP300 (100 nM; Enzo BML-SE451-0100) in assay buffer (25 mM HEPES pH 7.5, 150 mM NaCl, 2 mM DTT, 0.0001% Tween-20, 0.1 mg/m acetylated bovine serum albumin). This enzyme mix (3 μL) was added to white solid-bottom 1536-well plates (Greiner 789175) using a BioRaptr 2.0 Flying Reagent Dispenser (Let’s Go Robotics). Next, 20 nL of **1**-**3** (or DMSO equivalent) were transferred from stock solutions using a Kalypsys pintool and incubated with enzyme for 15 min. EP300 reactions were then initiated by addition of a 1 μL solution of acetyl-CoA (Sigma A2056) and histone H3 1-21 peptide substrate (AnaSpec AS-61702; ARTKQTARKSTGGKAPRKQLA-GGK(Biotin)-NH2). Following incubation for 1 h at room temperature, enzymatic activity was quenched and assessed by the addition of 4 μL of HTRF Detection Buffer (Cisbio 62SDBRDD) containing 50 μM anacardic Acid (Enzo 16611-84-0), Alexa Fluor 647-labeled anti-H3K9Ac antibody (Cell Signaling 4484) and Streptavidin Europium-Cryptate (Cisbio 610SAKLA). Plates were read using a PHERAstar FSX microplate reader with HTRF optical module (BMG LabTech) and normalized to no enzyme (negative) and DMSO (neutral) controls. Standard curves were generated through titration of peptide substrate and acetylated peptide [Lys(Ac)9]-Histone H3 (1-21; AnaSpec AS-64191) to benchmark activity. Acetyl-CoA and peptide substrates were varied for K_m_ determinations. Half-maximal inhibitor concentrations (Fig. 2) were determined at 20 nM histone H3 peptide (K_m_ = 18.3 nM) and the indicated concentration of acetyl-CoA (0.05 or 5 μM; K_m_ = 0.018 nM) except for **4** and **5**, whose IC_50_’s were determined at 50 nM H3 peptide and 200 μM acetyl-CoA. Replicate activity measurements in the presence of inhibitors were averaged and half-maximal inhibition values calculated from the nonlinear fit of dose-response data using GraphPad Prism 9.

**Cytotoxicity analyses**

EP300/CREBBP inhibitors **1**-**3** were analyzed for growth inhibition against the NCI-60 cell line panel using sulforhodamine B staining as previously described.^2-4^ To validate the NCI-60 relative activity trends using an orthogonal assay, we also assessed the activity of **1**-**3** against MCF-7 cells using an ATP-Glo assay. Briefly, MCF-7 cells were plated in white-walled 96-well plates (Corning 3610) at a density of 3 x 10^3^ cells/well and allowed to adhere for 24 h prior to treatment with compounds **1**-**3** for 96 h (0.25% maximum final concentrations of DMSO). Experiments were performed in quadruplicate and cell viability was determined using CellTiter-Glo Luminescent Cell Viability Assay (Promega G7572), according to the manufacturer’s instructions. Luminescence was recorded on a BioTek Synergy 2 plate reader and resulting data was reported as normalized percent cell viability, setting the average of DMSO vehicle control to 100%. The activity of **1**-**3** was also assessed in PANK4 KO models.^1^ Briefly, MCF10A cells were trypsinized and neutralized in serum free media lacking vitamin B5 (VB5-free media): custom DMEM lacking glucose, glutamine, and VB5 (US Biological Life Sciences) supplemented with 10% KnockOut Serum Replacement (Thermo Fisher Scientific/Gibco, 10828028), 10 µg/mL insulin, 0.5 mg/mL hydrocortisone, 20 ng/mL EGF, 100 ng/mL cholera toxin, 10 mM glucose, and 2 mM glutamine. Cells were centrifuged at 150 g for 3 min and washed in VB5-free media. Cells were again centrifuged at 150 g for 3 min and resuspended in VB5-free media. Cells were seeded at 2000 cells/well of a 96-well plate in VB5-free media supplemented with 1 µM VB5. After 24 h, three-fold drug dilution curves were made up in VB5-free media supplemented with 1 µM VB5, starting at 10 µM A-485 and 1 µM CPI-1612. The media was aspirated and replaced with drug media. 72 h after drugging, cells were fixed by addition of trichloroacetic acid to the media for a final concentration of 8.33% (w/v). Cell growth was measured by total protein staining via sulforhodamine B assay^3^ and expressed relative to vehicle-treated cells. Quadruplicate measurements were averaged and half-maximal inhibition values were calculated from the nonlinear fit of dose-response data in GraphPad Prism 9.

**Histone modification analyses**

***Treatment with EP300/CREBBP inhibitors***

MCF-7 cells were plated in 1700 µL culture medium at 1 x 10^6^ cells per well in 6-well dishes. After 16-24 h cells were treated with 300 µL of **1**-**3**-containing media to yield a final concentration of 8 nM, 40 nM, 200 nM, 1 µM, or 5 µM EP300/CREBBP inhibitor (0.1% final concentrations of DMSO). Controls were treated with vehicle DMSO (0.1%). After 3 h, MCF-7 cells were washed with ice-cold 1X PBS and scraped in 150 µL of Nuclear Isolation Buffer (NIB)^5^ (15 mM pH 7.5 Tris-HCl, 60 mM KCl, 15 mM NaCl, 5 mM MgCl_2_, 1 mM CaCl_2_, 250 mM sucrose, 1X protease inhibitor cocktail [Cell Signaling Technology #5871]), 1 mM DTT, and 10 mM sodium butyrate) with 0.1% IGEPAL CA-630 to lyse cells. Suspensions were transferred into tubes on ice and incubated for ≥5 min. Nuclei were pelleted (600 rcf, 4 °C , 5 min) and the supernatant discarded. Pelleted nuclei were subjected to two cycles of a wash with 150 µL NIB (without IGEPAL CA-630) followed by centrifugation (600 rcf, 4 °C , 5 min) in order to remove all IGEPAL CA-630. Nuclei were re-suspended in 400 µL 0.4 N H_2_SO_4_ and rotated at 4 °C for 16 h. The following day, samples were centrifuged (11,000 rcf, 4 °C , 10 min), supernatants transferred into new tubes (pellets discarded). 100 µL of 100% TCA was added to each sample (final TCA concentration: 20%) and tubes were inverted once. Histones were precipitated on ice for 2-6 h, then centrifuged (11,000 rcf, 4 °C , 5 min) and supernatants carefully discarded, with histones now visible as films on the sides and at the bottom of tubes. Histones were subjected to two cycles of washing (1 mL of ice-cold acetone + 1% 1M HCl, followed by 1 mL of 100% ice-cold acetone) followed by centrifugation (11,000 rcf, 4 °C , 5 min). Samples were air dried at room temperature and resuspended in 50 µL of ddH_2_O.

MCF10A cells (PANK4 KO+EV, KO+PANK4, KO + PANK4 mutant)^1^ were plated at 1.8 x 10^5^ cells per well in 10 cm dishes. After 16-24 h, media was aspirated and cells were treated with fresh media containing 40 nM, 200 nM, 1 µM, or 5 µM final concentrations of **1**-**3** (0.1% final concentrations of DMSO). Controls were treated with vehicle DMSO (0.1%). After 3 h, MCF10A cells were washed with 5 mL ice cold 1x PBS, scraped in 5 mL of 1x PBS, transferred to a pre-chilled 15 mL tube, centrifuged (800 rcf x 5 min), and snap frozen. Cell pellets were thawed on ice and 250 µL NIB^5^ (15 mM pH 7.5 Tris-HCl, 60 mM KCl, 15 mM NaCl, 5 mM MgCl_2_, 1 mM CaCl_2_, 250 mM sucrose, 1X protease inhibitor cocktail [Cell Signaling Technology #5871]), 1 mM DTT, and 10 mM sodium butyrate) with 0.3% IGEPAL CA-630 was added to lyse cells. Suspensions were transferred into tubes on ice and incubated for ≥5 min. Nuclei were pelleted (600 rcf, 4 °C , 5 min) and the supernatant discarded. Pelleted nuclei were subjected to three cycles of a wash with 250 µL NIB (without IGEPAL CA-630) followed by centrifugation (700 rcf, 4 °C , 5 min) in order to remove all IGEPAL CA-630. Nuclei were re-suspended in 250 µL 0.2 N H_2_SO_4_ and rotated at 4 °C for 2 h. Samples were then centrifuged (3,400 rcf, 4 °C , 5 min), supernatants transferred into new tubes (pellets discarded). 83 µL of 100% TCA was added to each sample (final TCA concentration: 20%) and tubes were inverted once. Histones were precipitated on ice for 16 h, then centrifuged (3,400 rcf, 4 °C , 5 min) and supernatants carefully discarded, with histones now visible as films on the sides and at the bottom of tubes. Histones were subjected to two cycles of washing (0.5 mL of ice-cold acetone + 1% 1M HCl, followed by 1 mL of 100% ice-cold acetone) followed by centrifugation (3,400 rcf, 4 °C , 5 min). Samples were air dried at room temperature and resuspended in 30 µL of ddH_2_O.

***Western blotting***

Histone samples were quantified using the Precision Red Protein Assay (Cytoskeleton #ADV02). 2 µg of histones from each condition were loaded onto 4-12% Bis-Tris NuPAGE gels (Invitrogen #NP0322 and #NP0323), using XCell SureLock Mini-Cells (Invitrogen EI0002) and MES running buffer (Invitrogen #NP0002) following manufacturer’s protocols. BenchMark Pre-stained Protein Ladder was used for all gels (Invitrogen 10748010). Separate gels were run for each antibody probed. SDS-PAGE gels were transferred to 0.2 µM pore nitrocellulose membranes (Novex, Life Technologies # LC2000) by either wet transfer electroblotting (Fig. 2b, 4e-f) at 30 volts for 1 h using a XCell II Blot Module (Invitrogen EI9051) and NuPAGE transfer buffer (Invitrogen NP00061) following the manufacturer’s protocols, or via iBlot dry transfers (Fig. 5f) using the iBlot dry blotting system (Invitrogen IB1001) and nitrocellulose transfer stacks (Invitrogen IB301001) at 20 volts for 8 min following the manufacturer’s protocols. Total protein content on Western blots was visualized by Ponceau staining followed by washing of membranes at least three times with 5% acetic acid in ddH_2_O. To visualize histone acetylation, membranes were blocked using StartingBlock (PBS) Blocking Buffer (Thermo Scientific 37538) for 20 min at room temperature and then incubated overnight at 4 °C in a solution containing the primary antibody at the indicated dilution. H3K18Ac (07-354) and H3K23Ac (07-355) were purchased from MilliporeSigma and used at 1:10,000 dilutions. H3K27Ac (8173S) antibody was purchased from Cell Signaling Technology and used at a 1:1,000 dilution. Membranes were washed the next day with 1x TBST at least 3 times and incubated for 1 h with anti-rabbit IgG HRP-linked antibody (Cell Signaling Technology 7074S) diluted to 1:1,000 in 1x TBST + 5% non-fat dry milk. Membranes were washed at least 3 times with 1x TBST, prior to imaging with Lumiglo (Cell Signaling Technology #7003) or SuperSignal ELISA Femto Substrate (Thermo Scientific 37074) according to manufacturer’s protocols. Colorimetric and chemiluminescent signals were detected using an Amersham ImageQuant 800 (Cytiva 29399482).

***LC-MS analyses of histone modifications***

T-47D cells were plated at 4 x 10^6^ cells per plate in 10 cm dishes in 10 mL of media. After 16-24 h cells were treated with 5, 50, or 500 nM of **3**. Compound **3** was dosed from stock solutions equal to 100x the final concentration in RPMI-1640. When necessary additional DMSO was added to maintain a constant concentration of 0.1%. Controls were treated with vehicle DMSO (0.1%). After 24 h, cells were washed with 5 mL ice cold 1x PBS, scraped in 5 mL of 1x PBS, transferred to a pre-chilled 15 mL tube, centrifuged (800 rcf x 5 min), and either snap frozen or immediately processed via the following protocol.^5^ Pellets were washed twice and resuspended in 400 µL NIB (15 mM pH 7.5 Tris-HCl, 60 mM KCl, 15 mM NaCl, 5 mM MgCl_2_, 1 mM CaCl_2_, 250 mM sucrose, 1X protease inhibitor cocktail [Cell Signaling Technology #5871]), 1 mM DTT, and 10 mM sodium butyrate) with 0.3% IGEPAL CA-630 to lyse cells. Suspensions were transferred into tubes on ice and incubated for ≥5 min. Nuclei were pelleted (700 rcf, 4 °C , 5 min) and the supernatant discarded. Pelleted nuclei were subjected to two cycles of a wash with 500 µL NIB (without IGEPAL CA-630) followed by centrifugation (700 rcf, 4 °C , 5 min) in order to remove all IGEPAL CA-630. Nuclei were re-suspended in 200 µL 0.2 N H_2_SO_4_ and rotated at 4 °C for 2 h. Samples were then centrifuged (3,400 rcf, 4 °C , 5 min) and supernatants transferred into new tubes (pellets discarded). 66 µL of 100% TCA was added to each sample (final TCA concentration: 20%) and tubes were inverted once. Histones were precipitated at 4 °C for 16 h, then centrifuged (3,400 rcf, 4 °C , 5 min) and supernatants carefully discarded, with histones now visible as films on the sides and at the bottom of tubes. Histones were subjected to two cycles of washing (0.5 mL of ice-cold acetone + 1% 1M HCl, followed by 1 mL of 100% ice-cold acetone) followed by centrifugation (11,000 rcf, 4 °C , 5 min) after each step. Samples were air dried at room temperature and resuspended in 30 µL of ddH_2_O. Histones were subjected to 2 min in a sonicating water bath, and centrifuged (3,400 rcf x 5 min) to remove any insoluble debris. Samples were analyzed by SDS-PAGE (Coomassie) and Western blotting to ensure clean extraction prior to LC-MS analysis.

Bottom-up LC-MS analysis of histone modifications was performed as reported previously.^5^ Briefly, 20 µg of purified histones were derivatized using propionic anhydride, followed by overnight digestion with 1 µg trypsin, and a second round of propionic anhydride derivatization to cap N-termini. The desalted peptides were then separated in a Thermo Scientific Acclaim PepMap 100 C18 HPLC Column (250mm length, 0.075mm I.D., Reversed Phase, 3 µm particle size) fitted on an Vanquish™ Neo UHPLC System (Thermo Scientific, San Jose, Ca, USA) using the HPLC gradient: 2% to 28% solvent B (A = 0.1% formic acid; B = 95% MeCN, 0.1% formic acid) over 45 min, from 28% to 80% solvent B in 5 min, 80% B for 10 min, all at a flow-rate of 300 nL/min. The samples were eluted into a QExactive-Orbitrap mass spectrometer (Thermo Scientific) and data acquired in a data-independent acquisition (DIA) as described previously.^5^ Accordingly, full scan MS (m/z 300−1100) was acquired in the Orbitrap with a resolution of 70,000 and an AGC target of 1 x 10^6^, and MS/MS was performed in centroid mode in the ion trap with sequential isolation windows of 24 m/z with an AGC target of 2 x 10^5^, a CID collision energy of 30 and a maximum injection time of 50 msec. An in-house software, EpiProfile was utilized to analyze the data.^6^ In this approach, the determination of chromatographic profile and discrimination of isobaric forms of peptides were determined using precursor and fragment extracted ions. The results were output as peptide relative ratios (% of total area under the extracted ion chromatogram of particular peptide form/sum of unmodified and modified forms belonging to the same peptide same amino acid sequence).

**In vitro and cellular analysis of photocaged inhibitors**

***HPLC analysis of photocaged inhibitor release***

To assess the ability of **4** to undergo photodeprotection to form **5** in the presence of UV light, we analyzed model reactions by HPLC. Briefly, 100 µM of **4** was dissolved in PBS (1% final concentration of DMSO) and irradiated in a UV-star 96 well plate for 0 min, 1 min, 2 min, or 10 min using 302 nm irradiation from a UVP Benchtop Variable Transilluminator (Analytik Jena P/N 95-0458-01, M-26V, 8-watt; high intensity setting). The irradiation source originated from below the plate and was conducted inside a UVP Multi-Doc-It Imaging System box (Analytik Jena P/N 97-0194-01) as a safety precaution. To assess conversion of **4** to **5**, irradiated reactions and controls were analyzed using an Agilent 1260 Infinity HPLC equipped with a Phenomenenex Kinetex C18 column (2.6 μm, 100 Angstrom, 100 × 4.6 mm inner diameter, H20-280322) and UV detector with monitoring at 280 nm. Solvents used were 0.1% TFA in H_2_O (A) and 0.1% TFA in acetonitrile (B), with a flow rate of 1 mL/min and 10 µL of sample injected. The method used 100% A from 0 to 2 min, followed by a linear gradient to 80/20 A/B from 2 to 10 min, 100% B from 10 to 12 min, and finally 100% A from 12 to 15 min. This method resulted in elution of **4** at approximately 7.9 min and **5** at approximately 6.1 min.

***Cellular analysis of photocaged inhibitor release***

MCF-7 cells were plated in 2000 µL culture medium at 5 x 10^5^ cells per well in 6-well dishes. After 16-24 h media was aspirated, 1700 µL fresh culture medium was added, and cells were treated with 300 µL of **4**-containing media to yield a final concentration of 10 µM, or 25 µM photocaged inhibitor (0.25% final concentrations of DMSO). Controls were treated with vehicle DMSO (0.25%). After 3 h, cells were washed once with PBS, 2 mL additional PBS was added, and cells were irradiated for 1 min with 302 nm, high intensity irradiation from below using the UVP Benchtop Variable Transilluminator. PBS was aspirated, media was replaced, and cells were returned to the incubator for 3 h before harvesting following the same histone extraction protocol for 6-well dishes described in the section *“Treatment with EP300/CREBBP inhibitors”*. Results are representative of two independent replicates.

**Supplementary schemes.**

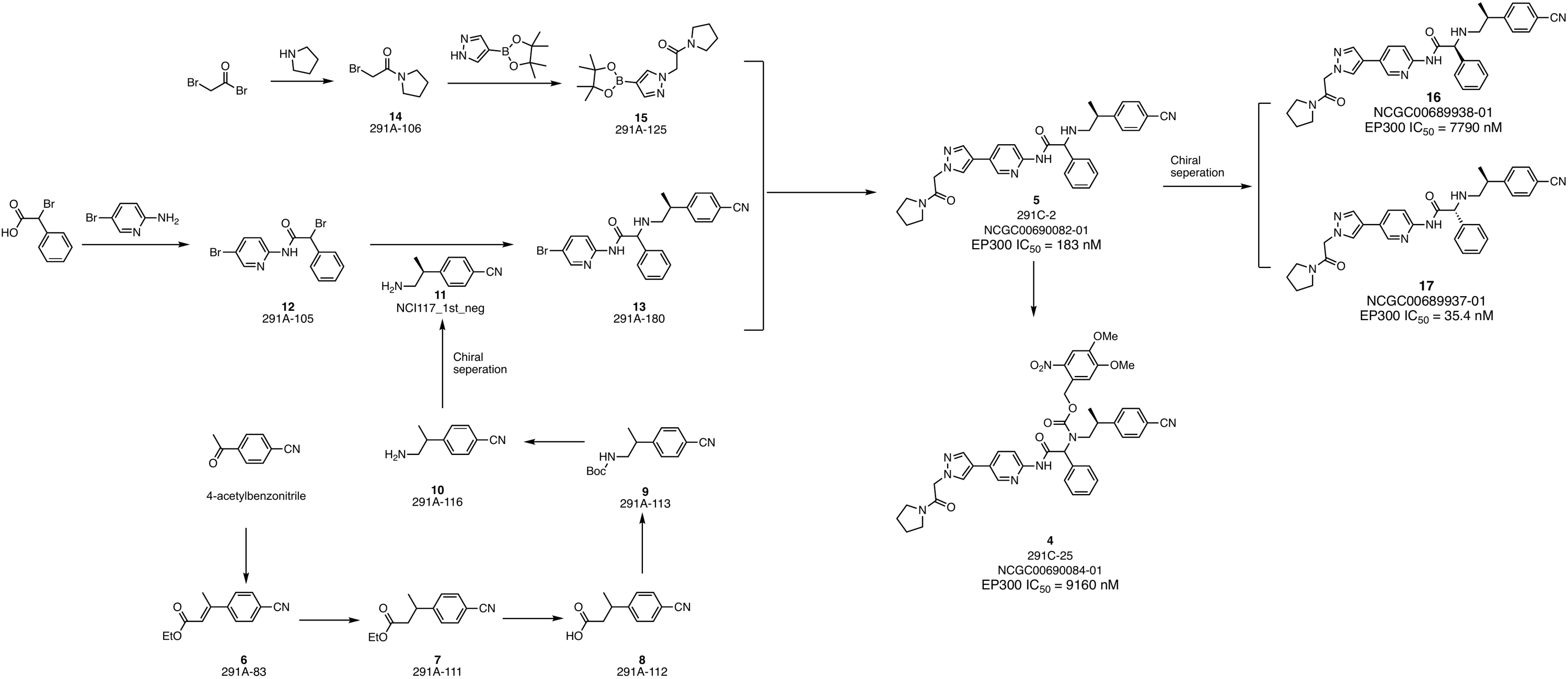

***Supplementary Scheme S1***. Synthesis of photocaged EP300 inhibitor.

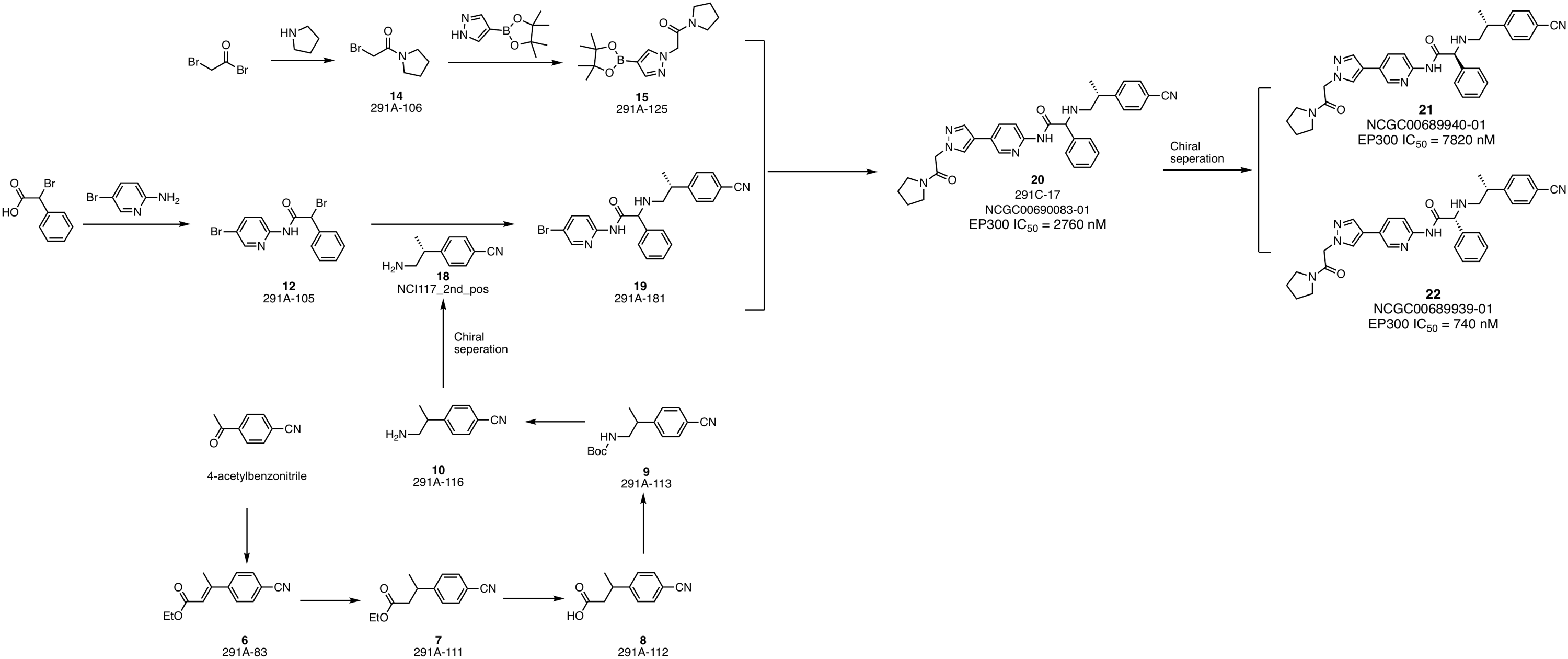

***Supplementary Scheme S2***. Synthesis of stereoisomers related to **5** used to verify biochemical potency and absolute configuration.

**Synthesis of photocaged inhibitor 4**

The precursor to photocaged inhibitor **4** was prepared according to the route published in the patent literature;^7^ However, with the hope of enabling future studies we also provide synthetic characterization data from our laboratory below.

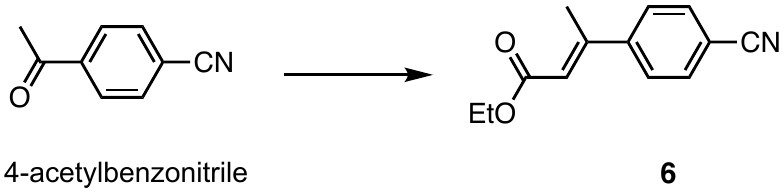

NaH (60% in mineral oil, 1.65 g, 68.88 mmol, 2 eq) was dissolved in dry THF, the mixture was cooled to 0 ^°C^, triethylphosphonoacetate (15.44 g, 68.88 mmol, 2 eq) was added dropwise to the solution before the resulting solution was stirred for 1 h at 0 °C. 4-acetylbenzonitrile (5 g, 34.44 mmol) was added to the reaction mixture, before the mixture was stirred for 1 h at 0 ^o^C and subsequently being gradually warmed to room temperature. The resulting mixture was stirred until full conversion before water and Et_2_O were added. The organic phase was separated, the aqueous phase was extracted with Et_2_O (3 x 20 mL), the combined organic phases were dried over MgSO_4_, concentrated in vacuo and purified by flash chromatography (1:5 EtOAc/hexane) to afford compound **6** as a colorless oil (6.6 g, 89%).^1^H NMR (400 MHz, CDCl_3_) δ 7.70 – 7.63 (m, 2H), 7.59 – 7.51 (m, 2H), 6.14 (q, *J* = 1.4 Hz, 1H), 4.23 (q, *J* = 7.1 Hz, 2H), 2.56 (d, *J* = 1.3 Hz, 3H), 1.32 (t, *J* = 7.1 Hz, 3H).

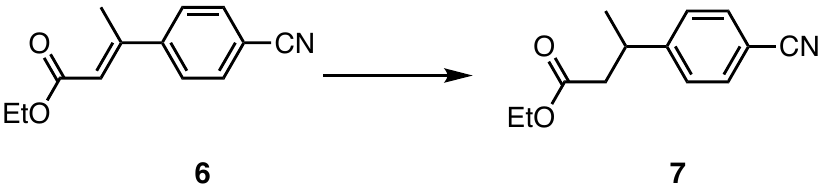

To a stirred solution of ethyl (*E*)-3-(4-cyanophenyl)but-2-enoate **6** (5.5 g, 25.55 mmol) in methanol/EtOAc (1:4, 140 mL) was added Pd/C (550 mg, 10% w/w, 50% moisture). The reaction was stirred at room temperature under an atmosphere of hydrogen gas for 3 h. The reaction mixture was diluted with EtOAc and filtered through a pad of celite. The combined organic layers were concentrated under reduced pressure to afford compound **7** (3.7 g, 67%). ^1^H NMR (400 MHz, CDCl_3_) δ 7.66 – 7.51 (m, 2H), 7.39 – 7.28 (m, 2H), 4.06 (qd, *J* = 7.2, 1.2 Hz, 2H), 3.34 (h, *J* = 7.2 Hz, 1H), 2.66 – 2.51 (m, 2H), 1.31 (d, *J* = 7.0 Hz, 3H), 1.17 (t, *J* = 7.1 Hz, 3H). ^13^C NMR (101 MHz, CDCl_3_) δ 171.67, 151.22, 132.39, 127.73, 118.93, 110.36, 60.52, 42.30, 36.62, 21.60, 14.15.

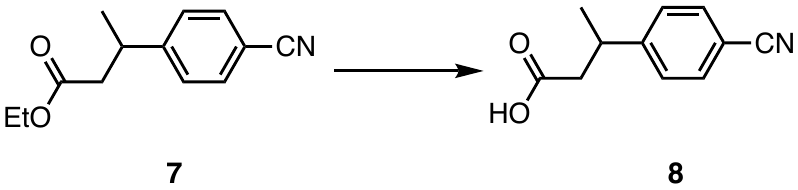

To a stirred solution of ethyl 3-(4-cyanophenyl)butanoate **7** (3.0 g, 13.81 mmol) in a mixture of MeOH : THF: H20 (4:2:1, 100 mL) was added LiOH (1.32 g, 55.23 mmol, 4 eq) at 5 ^o^C to 10 ^o^C. The resulting reaction mixture was stirred at room temperature for 1.5 h. After completion of reaction (monitored by TLC), the reaction solvent was evaporated. The residue was dissolved in water (10 mL) and extracted with EtOAc (2 x 15 mL). The pH of the aqueous layer adjusted to 3-4 with concentrated HCI. The precipitate that formed was filtered off to afford compound **8** (2.32 g, 89%) as a white solid. ^1^H NMR (500 MHz, CDCl_3_) δ 7.60 (d, *J* = 8.3 Hz, 2H), 7.38 – 7.29 (m, 2H), 3.32 (h, *J* = 7.2 Hz, 1H), 2.71 – 2.57 (m, 2H), 1.32 (d, *J* = 7.0 Hz, 3H). ^13^C NMR (126 MHz, CDCl_3_) δ 177.22, 150.79, 132.50, 127.68, 118.84, 110.53, 41.76, 36.25, 21.64.

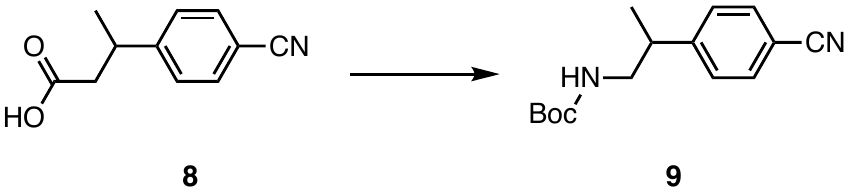

To a stirred solution of 3-(4-cyanophenyl)butanoic acid **8** (2.32 g, 12.26 mmol) in *tert*-butanol (65 mL) was added triethylamine (3.72 g, 36.78 mmol, 3.0 eq)at room temperature. Then the reaction mixture was cooled to 5-10 ^o^C and was added DPPA (5.74 g, 20.84 mmol, 1.7 eq) drop wise. After formation of acylazide, the reaction was stirred at 90 ^o^C overnight. The reaction mixture was diluted with water (40 mL) and extracted with EtOAc (2 x 40 mL). The combined organic layers were washed with brine, dried over Na_2_SO_4_ and concentrated under reduced pressure. The resulting residue was purified by flash chromatography (1:4 EtOAc/hexane) to afford compound **9** (2.04 g, 64%) as a colorless oil. ^1^H NMR (400 MHz, CDCl_3_) δ 7.67 – 7.54 (m, 2H), 7.31 (d, *J* = 8.0 Hz, 2H), 4.45 (s, 1H), 3.36 (dt, *J* = 13.1, 6.6 Hz, 1H), 3.21 (ddd, *J* = 13.8, 8.0, 5.9 Hz, 1H), 3.00 (p, *J* = 7.1 Hz, 1H), 1.40 (s, 9H), 1.27 (d, *J* = 6.9 Hz, 3H). ^13^C NMR (101 MHz, CDCl_3_) δ 155.92, 150.07, 132.54, 128.31, 119.03, 110.64, 79.63, 47.17, 40.61, 28.46, 18.78.

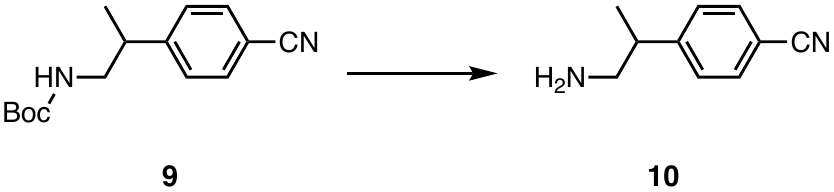

To a stirred solution *tert*-butyl (2-(4-cyanophenyl)propyl)carbamate **9** (2.0 g, 7.68 mmol) in 10 mL methanol was added a solution of 4M HCl in dioxane (10 mL) drop wise at 0 ^o^C. The resulting mixture was stirred at room temperature for 2 h. The reaction mixture was concentrated under reduced pressure to afford compound **10** (1.03 g, 84%) as a white solid. The chiral purification of **11** from racemic **10** is described in the next section. ^1^H NMR (400 MHz, *d6-*DMSO) δ 8.16 (s, 2H), 7.81 (d, *J* = 8.1 Hz, 2H), 7.53 (d, *J* = 8.0 Hz, 2H), 3.25 – 3.15 (m, 1H), 3.01 (p, *J* = 6.3, 5.8 Hz, 2H), 1.27 (d, *J* = 6.9 Hz, 3H). ^13^C NMR (101 MHz, *d6*-DMSO) δ 148.67, 132.57, 128.55, 118.85, 109.75, 44.27, 37.55, 19.00.

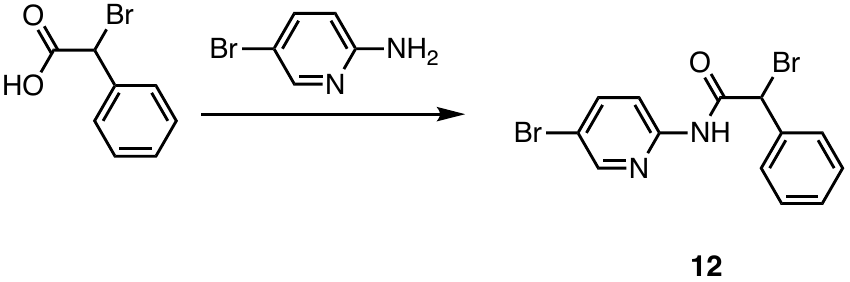

To a stirred solution of 5-bromopyridin-2-amine (1.5 g, 8.67 mmol) and 2-bromo-2-phenylacetic acid (2.05 g, 9.54 mmol, 1.1 eq) in 50 mL EtOAc was added n-propanephosphonic acid anhydride (T_3_P) (8.28 g, 13 mmol, 1.5 eq, 50% in EtOAc). The reaction mixture was stirred for 30 min at room temperature. Then DIPEA (2.24 g, 17.34 mmol, 2 eq) was added and the reaction mixture was heated at 60 ^°C^ for 3 h. The reaction mixture was poured into water and extracted with EtOAc (2 x 40 mL). The combined organic layers were washed with brine (20 mL), dried over Na_2_SO_4_ and concentrated under reduced pressure. The resulting residue was purified by flash chromatography (1:5 EtOAc/hexane) to afford compound **12** (520 mg, 16%) as a colorless oil. ^1^H NMR (400 MHz, *d4*-MeOH) δ 8.38 (dd, *J* = 2.5, 0.8 Hz, 1H), 8.08 (d, *J* = 8.9 Hz, 1H), 7.92 (dd, *J* = 8.9, 2.5 Hz, 1H), 7.68 – 7.59 (m, 2H), 7.42 – 7.32 (m, 3H), 5.76 (s, 1H). ^13^C NMR (101 MHz, MeOD) δ 168.50, 151.66, 150.18, 141.90, 137.88, 130.22, 129.89, 129.75, 116.66, 116.16, 49.71.

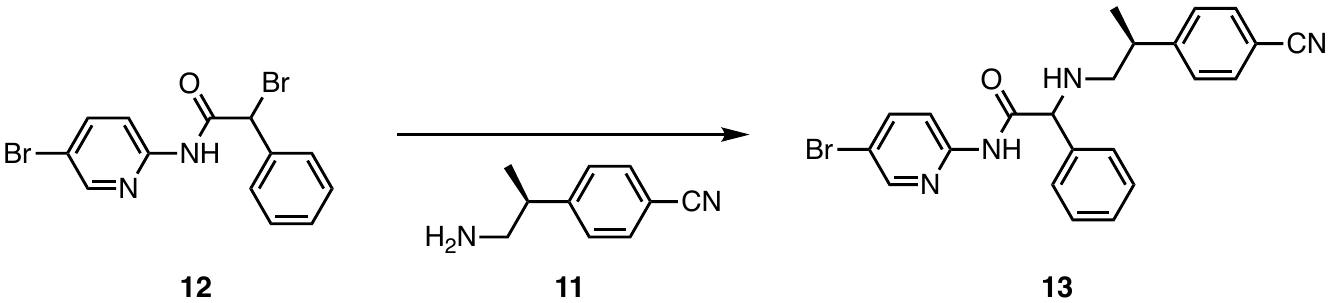

A mixture of 2-bromo-*N*-(5-bromopyridin-2-yl)-2-phenylacetamide **12** (720 mg, 1.95 mmol), (*R*)-4-(1-aminopropan-2-yl)benzonitrile **11** (374.08 mg, 2.33 mmol, 1.2 eq), and triethylamine (236.27 mg, 2.33 mmol, 1.2 eq) in 20 mL DMF was heated for 2 h at 60 ^°C^. After completion of the reaction, the reaction mixture was poured into ice cold water and extracted with EtOAc (2 x 40 mL). The combined organic layers were washed with brine, dried over Na_2_SO_4_ and concentrated under reduced pressure. The resulting residue was purified by flash chromatography (1:2 EtOAc/hexane) to afford compound **13** (710 mg, 81%) as a white solid. ^1^H NMR (500 MHz, MeOD) δ 8.35 (d, *J* = 2.5 Hz, 1H), 8.02 (dd, *J* = 19.0, 8.8 Hz, 1H), 7.88 (ddd, *J* = 8.8, 7.3, 2.5 Hz, 1H), 7.64 (d, *J* = 8.0 Hz, 2H), 7.52 – 7.38 (m, 2H), 7.38 – 7.14 (m, 5H), 4.33 (d, *J* = 16.8 Hz, 1H), 3.66 (s, 1H), 3.13 – 3.00 (m, 1H), 2.96 – 2.70 (m, 2H), 1.29 (dd, *J* = 14.9, 7.0 Hz, 3H). ^13^C NMR (126 MHz, MeOD) δ 173.49, 173.37, 152.71, 152.49, 151.29, 151.12, 150.04, 150.02, 141.94, 141.86, 139.88, 139.70, 133.60, 133.54, 129.87, 129.48, 129.38, 129.36, 128.54, 128.46, 120.05, 119.89, 116.35, 116.29, 115.83, 111.25, 111.22, 68.70, 68.37, 68.13, 56.27, 55.47, 41.73, 41.49, 19.94, 19.86.

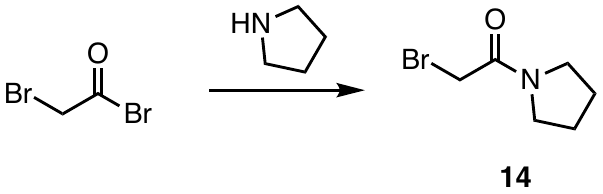

2-bromoacetyl bromide (6.39 g, 31.64 mmol, 1.5 eq) was added dropwise to a stirred solution of pyrrolidine (1.5 g, 21.09 mmol) and triethylamine (3.2 g, 31.64 mmol, 1.5 eq) in 20 mL DCM cooled to 0 ^°C^. The reaction was stirred at room temperature for 2 h. The reaction mixture was poured into cold 1*N* HCl solution (20 mL) and extracted with DCM (2 x 40 mL). The combined organic layers were washed with brine, dried over Na_2_SO_4_ and concentrated under reduced pressure. The resulting residue was purified by flash chromatography (1:1 EtOAc/hexane) to afford compound **14** (2.17 g, 54%) as a brown oil. ^1^H NMR (500 MHz, CDCl_3_) δ 3.80 (s, 2H), 3.50 (dt, *J* = 20.6, 6.9 Hz, 4H), 2.06 – 1.95 (m, 2H), 1.93 – 1.83 (m, 2H).

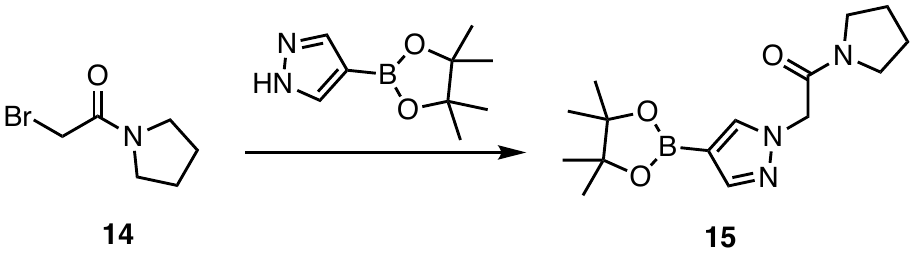

To a stirred solution of 4-(4,4,5,5-tetramethyl-1,3,2-dioxaborolan-2-yl)-1*H*-pyrazole **14** (700 mg, 3.61 mmol) in 10 mL dry DMF was added NaH (187.57 mg, 60%, 4.69 mmol, 1.3 eq) at 0 ^°C^. The reaction was stirred at room temperature for 15 min. To this 2-bromo-1-(pyrrolidin-1-yl)ethan-1-one (1.04 g, 5.41 mmol, 1.5 eq) was added at 0 ^°C^ and stirred for 30 min at same temperature. The reaction mixture was then stirred at room temperature for another 1 h. The reaction mixture was poured into ice cold water and extracted with DCM (2 x 40 mL). The combined organic layers were washed with brine, dried over Na_2_SO_4_ and concentrated under reduced pressure. The resulting residue was purified by flash chromatography (1:10 MeOH/DCM) to afford compound **15** (500 mg, 45%) as a colorless oil. ^1^H NMR (400 MHz, CDCl_3_) δ 7.80 (d, *J* = 19.2 Hz, 2H), 4.90 (s, 2H), 3.47 (dt, *J* = 11.2, 7.0 Hz, 4H), 2.02 – 1.92 (m, 2H), 1.90 – 1.80 (m, 2H), 1.30 (s, 12H). ^13^C NMR (101 MHz, CDCl_3_) δ 164.75, 145.83, 137.96, 83.39, 54.12, 46.39, 46.30, 26.30, 24.98, 24.92, 24.18.

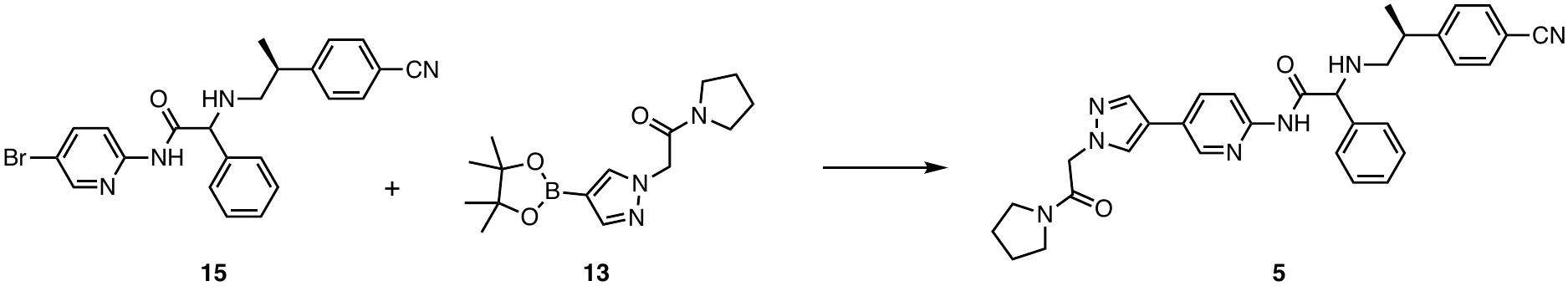

A mixture of *N*-(5-bromopyridin-2-yl)-2-(((*R/S*)-2-(4-cyanophenyl)propyl)amino)-2-phenylacetamide **15** (130 mg, 0.289 mmol), 1-(pyrrolidin-1-yl)-2-(4-(4,4,5,5-tetramethyl-1,3,2-dioxaborolan-2-yl)-1*H*-pyrazol-1-yl)ethan-1-one **13** (97.12 mg, 0.318 mmol, 1.1 eq), and cesium carbonate (282.78 mg, 0.868 mmol, 3 eq)in 4:1 dioxane:water (8 mL) was purged for 20 min with argon. S-Phos Pd-precatalyst G3 (22.57 mg, 0.029 mmol, 0.1 eq) was added and purging was continued for another 10 min. The reaction mixture was heated in a sealed tube at 95 ^°C^ for 2 h. Then the reaction mixture was treated with water and extracted with EtOAc (2 x 20 mL). The combined organic layers were washed with brine, dried over Na_2_SO_4_ and concentrated under reduced pressure. The resulting residue was purified by flash chromatography (1:10 MeOH/DCM) to afford compound **5** (70 mg, 47%) as a white solid. ^1^H NMR (500 MHz, MeOD) δ 8.52 (d, *J* = 2.4 Hz, 1H), 8.12 – 8.00 (m, 2H), 7.96 – 7.86 (m, 2H), 7.64 (dd, *J* = 8.3, 3.0 Hz, 2H), 7.56 – 7.11 (m, 8H), 5.08 (d, *J* = 1.6 Hz, 2H), 4.34 (d, *J* = 13.1 Hz, 1H), 3.68 (s, 1H), 3.59 (t, *J* = 6.9 Hz, 2H), 3.46 (t, *J* = 7.0 Hz, 2H), 3.13 – 3.02 (m, 1H), 2.97 – 2.74 (m, 2H), 2.04 (p, *J* = 6.9 Hz, 2H), 1.91 (p, *J* = 6.9 Hz, 2H), 1.34 – 1.27 (m, 3H), 1.20 (s, 9H).

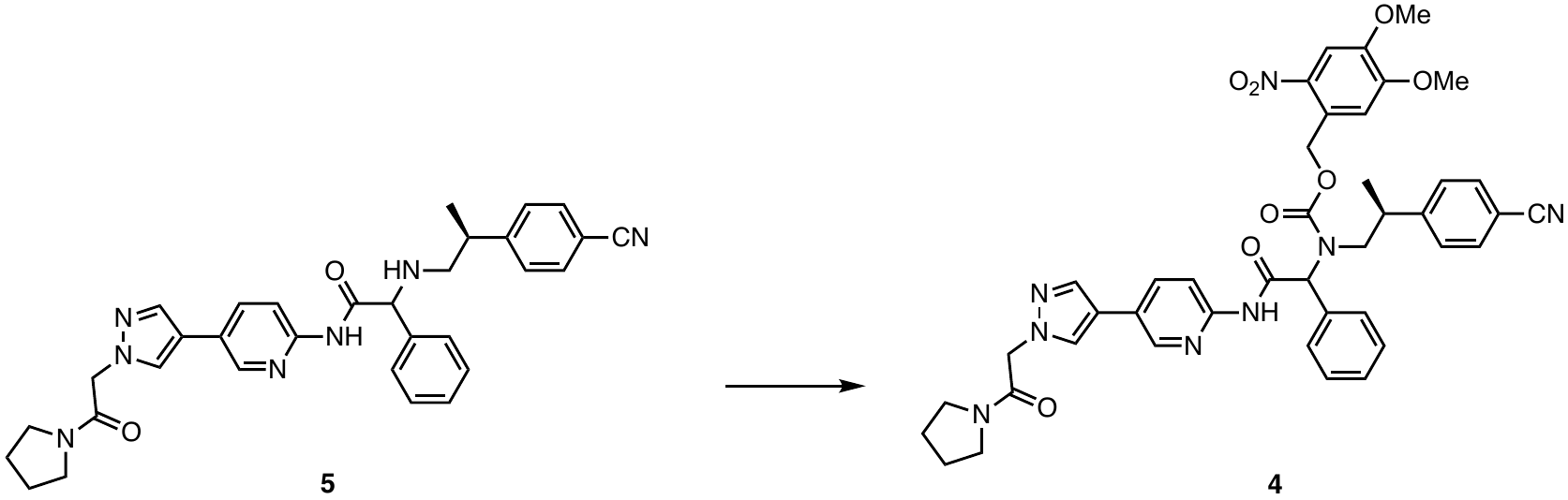

6-Nitroveratryl chloroformate (NVOC) (13.21 mg, 47.93 mmol, 1.05 eq) was added with stirring to a solution of 2-(((*R/S*)-2-(4-cyanophenyl)propyl)amino)-*N*-(5-(1-(2-oxo-2-(pyrrolidin-1-yl)ethyl)-1*H*-pyrazol-4-yl)pyridin-2-yl)-2-phenylacetamide **5** (25 mg, 45.65 mmol) and NaHCO_3_ (4.03 mg, 47.93 mmol, 1.05 equi) in CHCl_3_/Et_2_O (9:1, 10 mL) at 5 ^°C^. The solution was stirred for 2 h, before being treated with saturated aqueous NaHCO_3_ (1 x 10 mL) and H_2_O (2 x 10 mL). The organic phase was washed with brine, dried over Na_2_SO_4_ and concentrated under reduced pressure. The resulting residue was purified by flash chromatography (1:10 MeOH/DCM) to afford compound **4** (10 mg, 28%) as a white solid. ^1^H NMR (400 MHz, DMSO) δ 10.68 (s, 1H), 8.53 (s, 1H), 8.18 – 7.87 (m, 4H), 7.76 – 6.89 (m, 11H), 5.99 (d, *J* = 31.0 Hz, 1H), 5.53 – 5.21 (m, 2H), 5.02 (d, *J* = 12.7 Hz, 2H), 4.10 – 3.68 (m, 6H), 3.48 (dt, *J* = 14.4, 6.8 Hz, 3H), 3.40 – 3.24 (m, 5H), 1.91 (h, *J* = 6.5 Hz, 2H), 1.79 (p, *J* = 7.0 Hz, 2H), 1.11 – 0.70 (m, 4H). ^13^C NMR (101 MHz, DMSO) δ 169.51, 164.97, 164.72, 153.20, 150.22, 144.24, 138.48, 136.03, 134.64, 134.31, 132.12, 131.28, 130.06, 128.74, 128.53, 128.13, 127.91, 124.57, 118.87, 118.42, 113.44, 108.99, 108.11, 105.18, 73.50, 63.99, 56.18, 56.05, 53.66, 53.33, 45.74, 45.69, 45.18, 45.13, 29.01, 25.63, 24.94, 23.69, 17.60. MS (EI) (*m*/*z*): [M+H]^+^ calc’d for C_42_H_43_N_8_O_8_ predicted 787.3, obs’d 787.5

**Chiral separation and absolute configuration of 4 and 5 and related compounds**

***Development of chiral separation methods.***

The development and optimization of analytical chiral separation methods for chiral mixtures by normal phase liquid chromatography (LC) was performed on an Agilent 1200 HPLC system (Agilent Technologies, Santa Clara, CA, U.S.A.) equipped with a diode array detector (DAD), quaternary pump, multicolumn thermostat, autosampler, and a PDR-Chiral Advanced Laser Polarimeter (ALP) (PDR-Chiral, Inc., Lake Park, FL, U.S.A.). Samples were screened across 9 analytical chiral columns (CHIRALPAK AD, AS, IA, IC, ID, IF, and IG (Chiral Technologies, West Chester, PA, U.S.A.); CHIRALCEL OJ (Chiral Technologies, West Chester, PA, U.S.A.); (R,R) Whelk-O 1(Regis Technologies, Morton Grove, IL, U.S.A.) with 3 isocratic mobile phases (methanol (MeOH); hexane/ethanol (EtOH) (40:60 %v/v); hexane/isopropanol (iPrOH) (40:60 %v/v)) to provide separation results for 27 different methods. The enantiomers were identified by either a positive or negative peak in the ALP chromatogram. The preferred method for each sample was then further optimized through modification of the solvent system. This included the addition of 0.1% diethylamine (DEA) or 0.1% isopropylamine to hexane for the bi-solvent mobile phases or addition to MeOH for the single solvent mobile phase. The optimized methods were transferred to preparative separation by use of the corresponding preparative chiral column.

***Preparative chiral separation of 4-(1-aminopropan-2-yl)benzonitrile (10).***

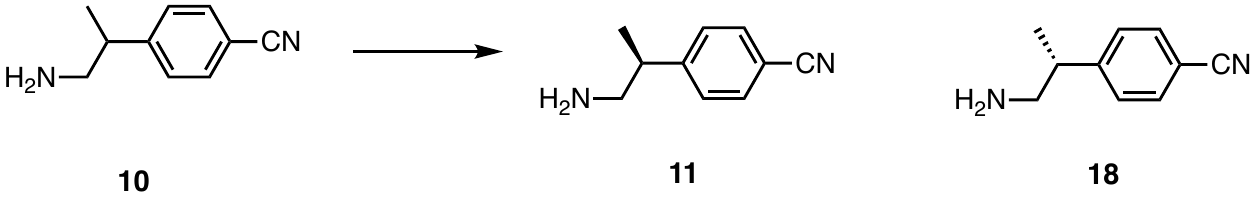

Preparative chiral separation of the racemic mixture (**10**) by normal phase liquid chromatography (LC) was performed on an Agilent 1200 HPLC system (Agilent Technologies, Santa Clara, CA, U.S.A.) equipped with a diode array detector (DAD), 1200 preparative pumps, a 5 mL sample loop, and a direct injection valve using a CHIRALPAK IG column (5 x 50 cm, particle size 20 µm) (Chiral Technologies, West Chester, PA, U.S.A.). The isocratic mobile phase used for the separation was hexane/ethanol/isopropylamine (40:60:0.04 %v/v) eluting at a flow rate of 40 mL/min with detection at 230 nm. A 4 mL aliquot at a concentration of 30 mg/mL in MeOH was injected onto the column for each run with a total of 1220 mg of crude racemic material being separated. The solvent of the collected fractions was removed using rotary evaporation to provide 471 mg of **11** and 460 mg of **18**. The optical activity of the enantiomers was confirmed using the Agilent 1200 LC system configured for analytical chiral analysis with detection by the PDR-Chiral ALP (PDR-Chiral, Inc., Lake Park, FL, U.S.A.). The negative rotating enantiomer (**11**) eluted first (RT = 34-47 min) being isolated in 95.2% ee and the positive rotating enantiomer (**18**) eluted second (RT = 41-36 min) being isolated in 98.7% ee.

***Preparative chiral separation of 5***

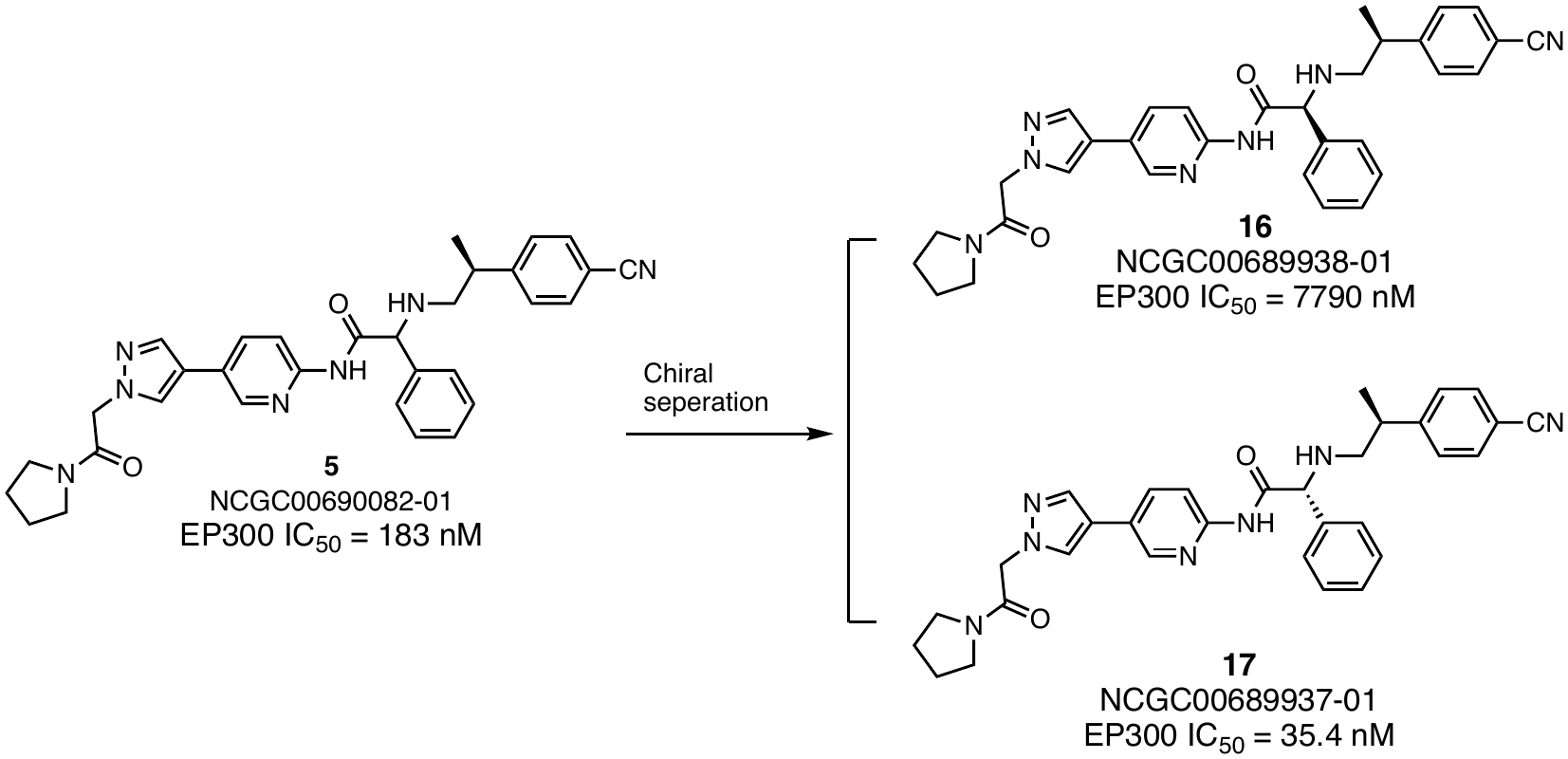

Preparative chiral separation of the diastereomeric mixture (**5**) by normal phase liquid chromatography (LC) was performed on an Agilent 1200 HPLC system (Agilent Technologies, Santa Clara, CA, U.S.A.) equipped with a diode array detector (DAD), 1200 preparative pumps, a 5 mL sample loop, and a direct injection valve using a (R,R) Whelk-O 1 column (5 x 50 cm, particle size 16 µm) (Regis Technologies, Morton Grove, IL, U.S.A.). The isocratic mobile phase used for the separation was MeOH/DEA (100:0.1 %v/v) eluting at a flow rate of 40 mL/min with detection at 230 nm. A 1 mL aliquot of **5** at a concentration of 28 mg/mL in MeOH was injected onto the column for each run with a total of 28 mg of crude racemic material being separated. The solvent of the collected fractions was removed using rotary evaporation to provide 8.5 mg of **16** (NCGC00689938-01) and 6.8 mg of **17** (NCGC00689937-01). The optical activity of the enantiomers was confirmed using the Agilent 1200 LC system configured for analytical chiral analysis with detection by the PDR-Chiral ALP (PDR-Chiral, Inc., Lake Park, FL, U.S.A.). The positive rotating enantiomer (**16**) eluted first (RT = 70-76 min) being isolated in >99% ee and the negative rotating enantiomer (**17**) eluted second (RT = 114-124 min) being isolated in >99% ee.

***Preparative chiral separation of 20.***

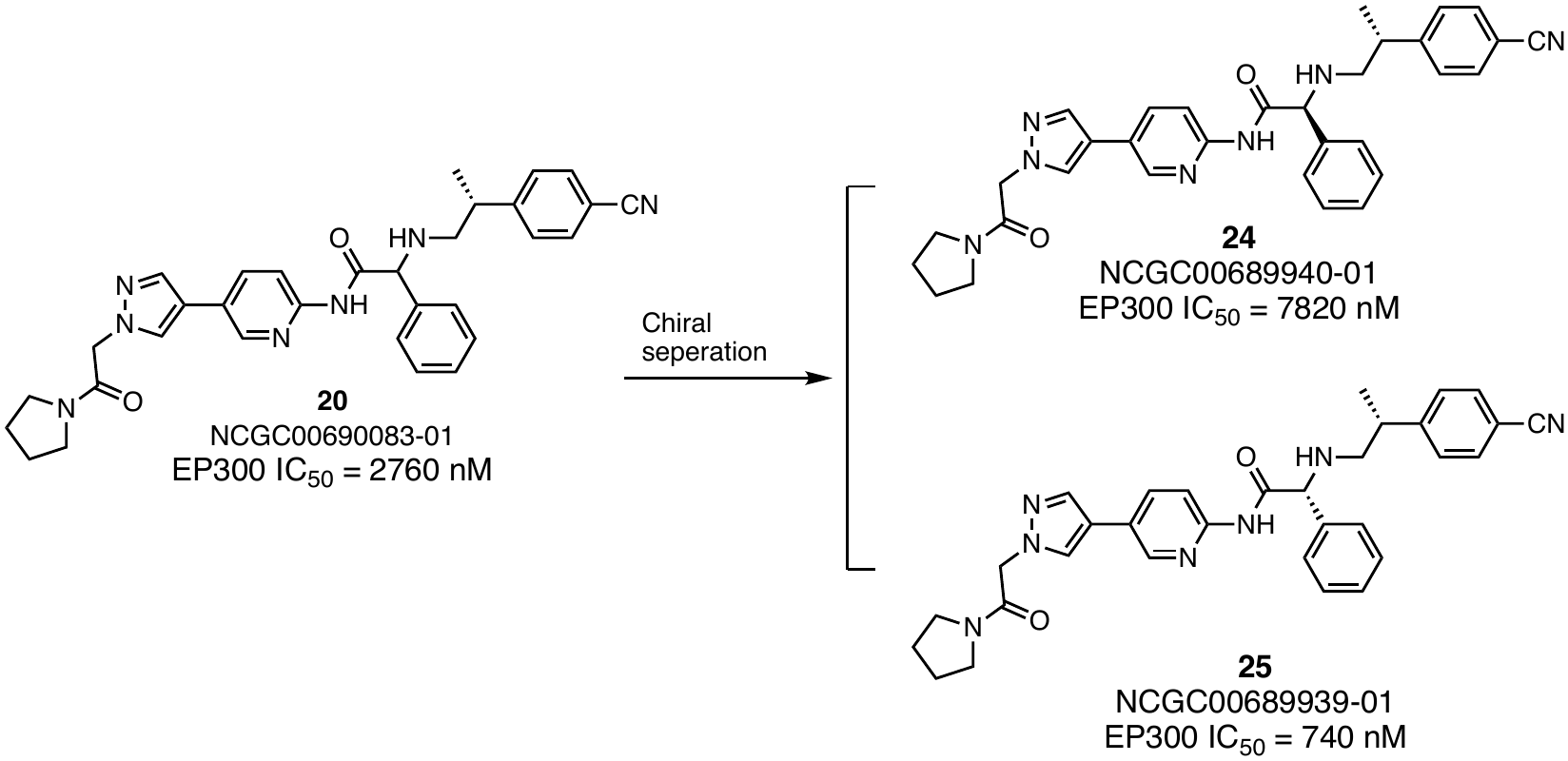

Preparative chiral separation of the diastereomeric mixture (**20**) by normal phase liquid chromatography (LC) was performed on an Agilent 1200 HPLC system (Agilent Technologies, Santa Clara, CA, U.S.A.) equipped with a diode array detector (DAD), 1200 preparative pumps, a 5 mL sample loop, and a direct injection valve using a (R,R) Whelk-O 1 column (5 x 50 cm, particle size 16 µm) (Regis Technologies, Morton Grove, IL, U.S.A.). The isocratic mobile phase used for the separation was MeOH/DEA (100:0.1 %v/v) eluting at a flow rate of 40 mL/min with detection at 230 nm. A 1 mL aliquot of **20** at a concentration of 30 mg/mL in MeOH was injected onto the column for each run with a total of 30 mg of crude racemic material being separated. The solvent of the collected fractions was removed using rotary evaporation to provide 5.6 mg of **24** (NCGC00689940-01) and 5.8 mg of **25** (NCGC00689939-01). The optical activity of the enantiomers was confirmed using the Agilent 1200 LC system configured for analytical chiral analysis with detection by the PDR-Chiral ALP (PDR-Chiral, Inc., Lake Park, FL, U.S.A.). The positive rotating enantiomer (**24**) eluted first (RT = 65-72 min) being isolated in >99% ee and the negative rotating enantiomer (**25**) eluted second (RT = 111-122 min) being isolated in >99% ee.

***Determination of absolute configuration by vibrational circular dichroism (VCD) analysis.***

VCD measurements were performed on a BioTools ChiralIR-2X with DualPEM FT-VCD instrument (BioTools, Inc., Jupiter, FL, U.S.A.) with a resolution of 4 cm^−1^ and a PEM focus frequency of 1400 cm^−1^. A 7 mg aliquot of either **11** or **18** was dissolved in 1250 µL of CDCl_3_ and then pipetted into a BaF_2_ IR cell with a path length of 100 µm. Each enantiomer was measured for 22 h followed by solvent subtraction of the IR spectra and correction of the VCD spectra with the half-difference method. The VCD spectra were obtained and processed in the same manner for **17** (NCGC00689937-01), **16** (NCGC00689938-01), **25** (NCGC00689939-01), and **24** (NCGC00689940-01) at a concentration of 1-2 mg/80-90 µL CDCl_3_.

A GMMX (MMFF94) search of the *R*-enantiomer of **11** found 11 low-energy conformers that were minimized using Gaussian 09 at the cc-pVTZ/B3LYP level with the CPCM solvent (CDCl_3_) model. IR and VCD frequencies were calculated at the same level, Boltzmann averaged, and plotted with a line width of 6 cm^−1^. IR and VCD spectra were frequency scaled by a factor of 0.976 and compared to experimental data. Comparison of the calculated and observed spectra found an IR similarity of 72.1, a VCD similarity of 73.8, and an enantiomeric similarity index (ESI) of 62.3. Based on these results, it was determined that **11** has an absolute configuration of R and **18** has an absolute configuration of S with a 99% confidence level.

The minimal sample amounts (~5 mg) available for the 4 diastereomers (**16**, **17**, **24**, **25**) provided low intensity IR spectra for **16** (NCGC00689938-01) and **17** (NCGC00689937-01), which generated VCD spectra with small signal size resulting in an overall confidence level of 75% for the VCD spectra. Comparing the frequencies and patterns in the IR spectra, it was determined that **16** (NCGC00689938-01) and **25** (NCGC00689939-01) are enantiomers and **17** (NCGC00689937-01) and **24** (NCGC00689940-01) are enantiomers. A GMMX (MMFF94) search of all potential diastereomers found 558 low-energy conformers that were minimized using Gaussian 09 at the 6-31Gd/B3LYP level with the CPCM solvent (CDCl_3_) model. IR and VCD frequencies were calculated at the same level, Boltzmann averaged, and plotted with a line width of 6 cm^−1^. IR and VCD spectra were frequency scaled by a factor of 0.976 and compared to experimental data. Based on the calculations, the phenyl (Ph) epimers exhibited much larger differences in the VCD spectra than the methyl (Me) epimers. This observation was used to correlate the experimental VCD spectra based on the similarity of the Ph stereochemistry. As such, **17** (NCGC00689937-01) and **25** (NCGC00689939-01) were one grouping while **16** (NCGC00689938-01) and **24** (NCGC00689940-01) were another grouping. Comparing these two sets of experimental VCD spectra to the calculated VCD spectra showed **17** (NCGC00689937-01) and **25** (NCGC00689939-01) correlated best to the calculated spectra where the Ph stereocenter had an *R* configuration. Additionally, it was known **17** (NCGC00689937-01) and **16 (**NCGC00689938-01) had an *S*-Me configuration and **25** (NCGC00689939-01) and **24** (NCGC00689940-01) had an *R*-Me configuration conveyed from their respective starting materials, **11** and **18**. Using all this data together enabled the absolute configuration assignment of all compounds: **16** (NCGC00689938-01) is (*S*-Ph,*S*-Me); **17** (NCGC00689937-01) is (*R*-Ph,*S*-Me); **24** (NCGC00689940-01) is (*S*-Ph,*R*-Me); and **25** (NCGC00689939-01) is (*R*-Ph,*R*-Me). This is also consistent with the IC_50_ values generated for these compounds using EP300 biochemical inhibition assays, as the compound that matches the reported stereochemistry of CPI-1612^8^ **17** is the most potent of the series (**17**; IC_50_ = 37 nM), followed by **25** (IC_50_ = 740 nM), **24** (IC_50_ = 7820 nM), and **z** (IC_50_ = 7790 nM).

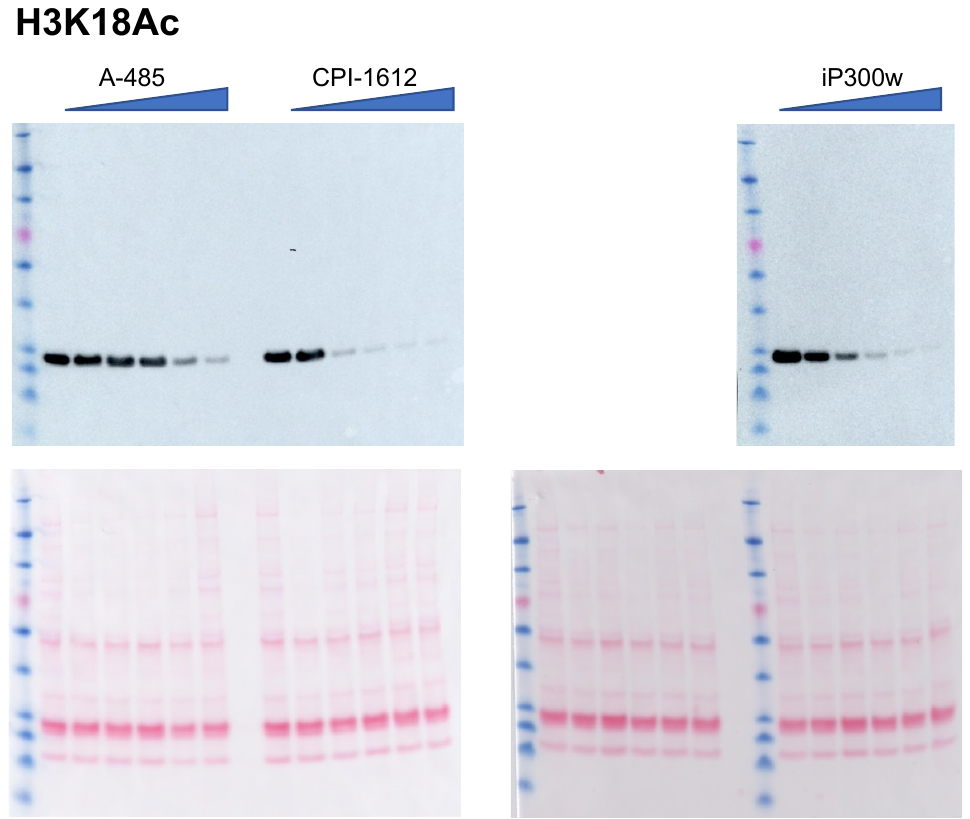
**Full Western blot images**

Full Western blots for Figure 3b, H3K18Ac mark. Top: anti-H3K18Ac. Bottom: Ponceau. Wet transfer method used; 10 s exposure time using Lumiglo imaging solution for both blots.

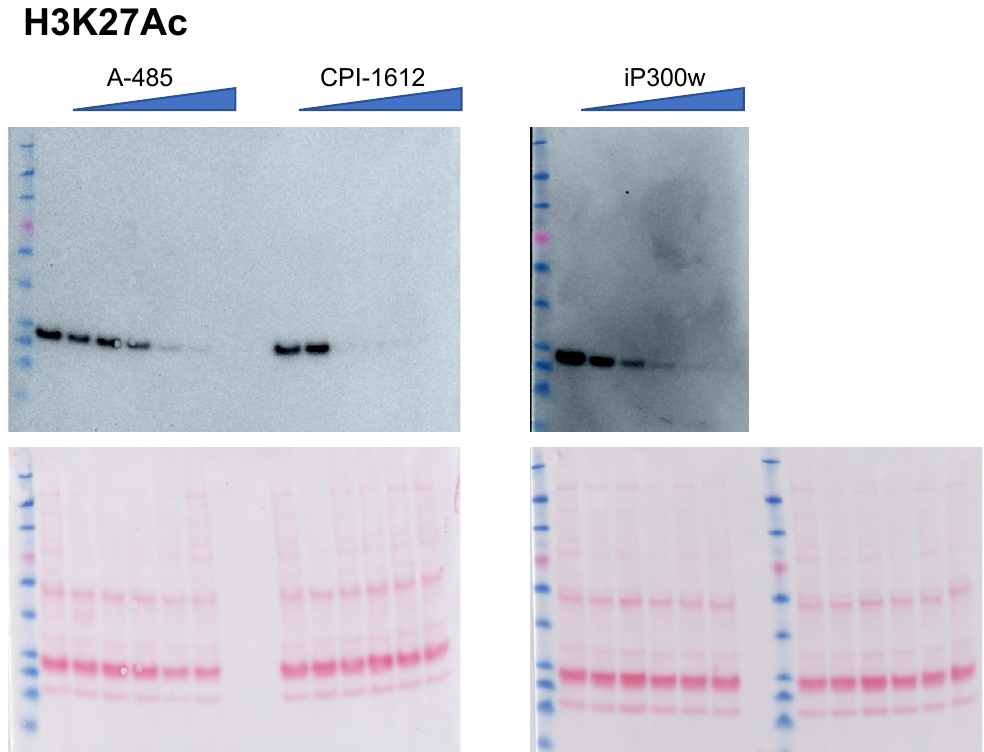

Full Western blots for Figure 3b, H3K27Ac mark. Top: anti-H3K27Ac. Bottom: Ponceau. Wet transfer method used; 1 min exposure time using Lumiglo imaging solution for both blots.

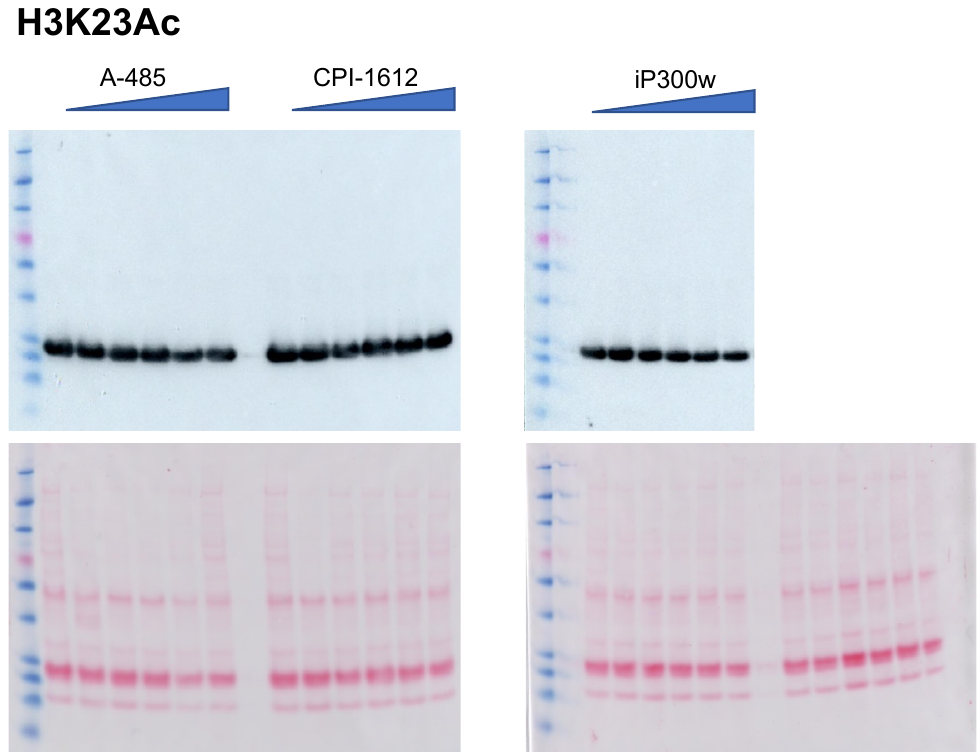

Full Western blots for Figure 3b, H3K23Ac mark. Top: anti-H3K18Ac. Bottom: Ponceau. Wet transfer method used; 5 s exposure time using Lumiglo imaging solution for both blots.

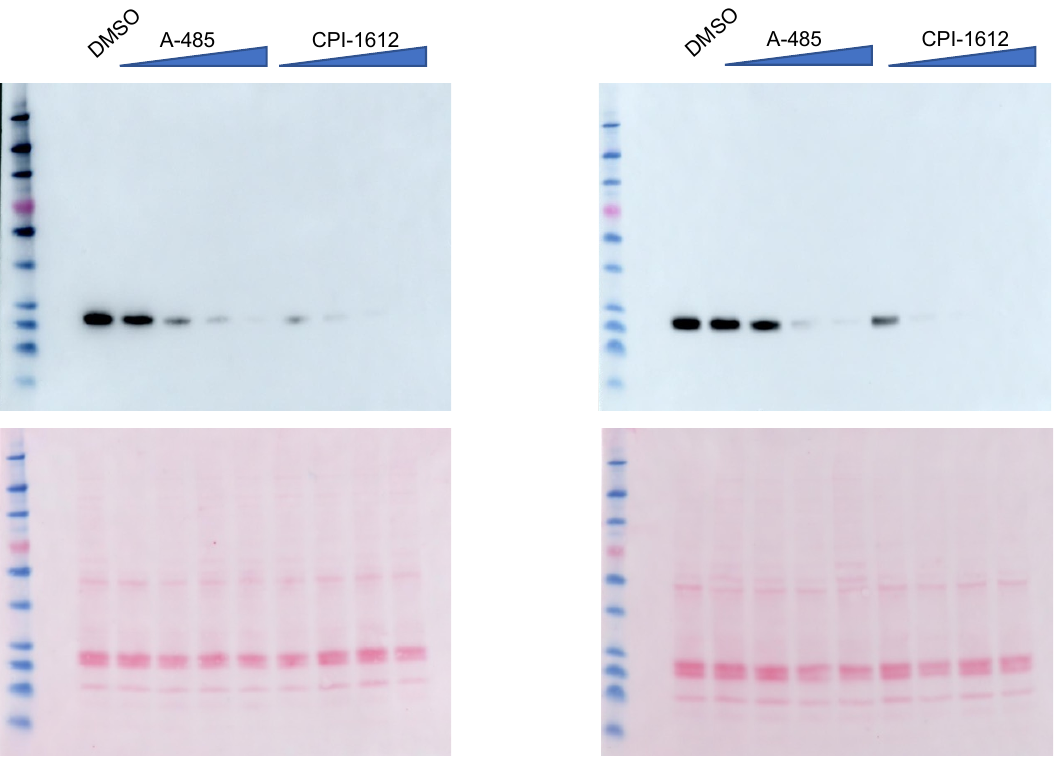

Full Western blots for Figure 4e. PANK4 WT samples on left blot and PANK4 KO on right blot. Top: anti-H3K18Ac. Bottom: Ponceau. Wet transfer method used; 2 s exposure time using SuperSignal imaging solution for both blots.

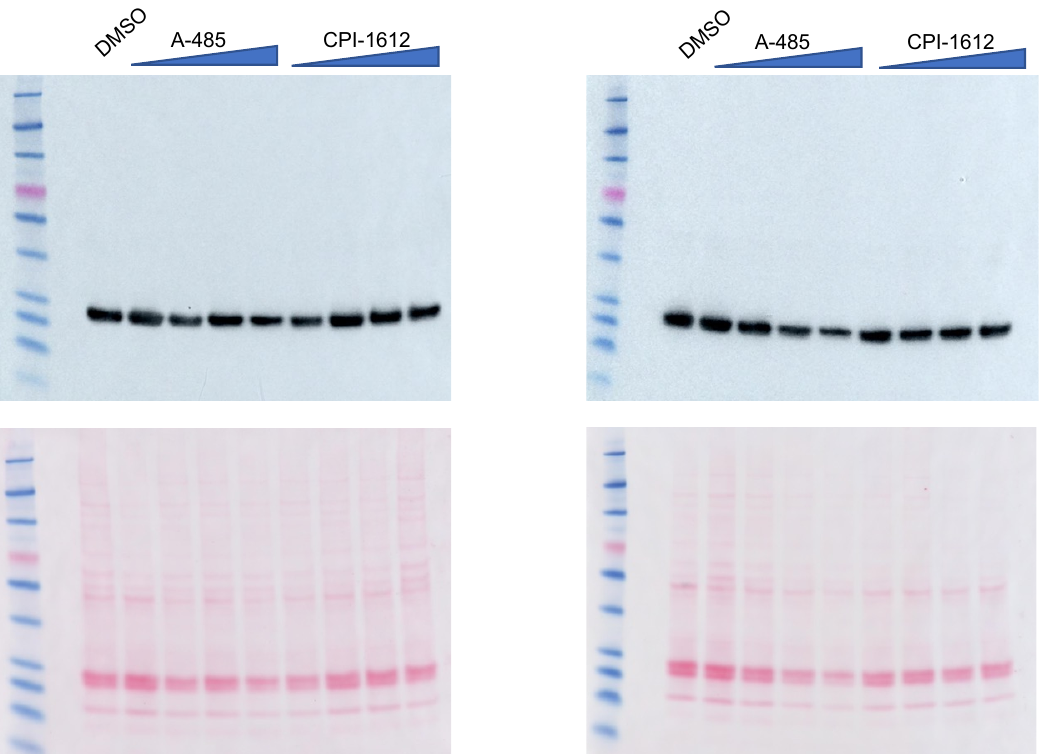

Full Western blots for Figure 4f. PANK4 WT samples on left blot and PANK4 KO on right blot. Top: anti-H3K23Ac. Bottom: Ponceau. Wet transfer method used; 2 s exposure time using Lumiglo imaging solution for both blots.

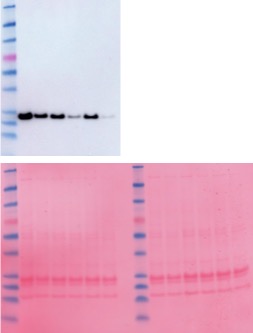

Full Western blots for Figure 5f. Top: anti-H3K18Ac. Bottom: Ponceau. iBlot transfer method used; 30 s exposure time using Lumiglo imaging solution for both blots.
